# Rate variation in the evolution of non-coding DNA associated with social evolution in bees

**DOI:** 10.1101/461079

**Authors:** Benjamin E.R. Rubin, Beryl M. Jones, Brendan G. Hunt, Sarah D. Kocher

## Abstract

The evolutionary origins of eusociality represent increases in complexity from individual to caste-based, group reproduction. These behavioral transitions have been hypothesized to go hand-in-hand with an increased ability to regulate when and where genes are expressed. Bees have convergently evolved eusociality up to five times, providing a framework to test this hypothesis. To examine potential links between putative gene regulatory elements and social evolution, we compare alignable, non-coding sequences in eleven diverse bee species, encompassing three independent origins of reproductive division of labor and two elaborations of eusocial complexity. We find that rates of evolution in a number of non-coding sequences correlate with key social transitions in bees. Interestingly, while we find little evidence for convergent rate changes associated with independent origins of social behavior, a number of molecular pathways exhibit convergent rate changes in conjunction with subsequent elaborations of social organization. We also present evidence that many novel non-coding regions may have been recruited alongside the origin of sociality in corbiculate bees; these loci could represent gene regulatory elements associated with division of labor within this group. Thus, our findings are consistent with the hypothesis that gene regulatory innovations are associated with the evolution of eusociality and illustrate how a thorough examination of both coding and non-coding sequence can provide a more complete understanding of the molecular mechanisms underlying behavioral evolution.

## Introduction

Many genomic sequences that do not encode proteins play essential roles in gene regulation across animals [1] and plants [2]. The breadth of knowledge of these non-coding regulatory elements has been built primarily upon the large number of plant and vertebrate genomes that have been sequenced over the past decade. However, the high degree of conservation that exists in a subset of these non-coding regions [1,3] means that comparative methods can be used to identify similar non-coding elements even in the recently sequenced genomes of non-model taxa that frequently lack the resources needed to characterize regulatory regions via functional assays. Insects, in particular, have been the focus of many *de novo* genome sequencing projects yet, outside of *Drosophila* [4], the non-coding regulatory landscape of these taxa has been the target of few studies.

Here, we take advantage of eleven publicly-available bee genomes [5–9] to examine how noncoding elements change as eusociality – an extreme form of social behavior found primarily in insects and mammals – has convergently evolved and increased in complexity within bees. Eusociality is of particular interest to evolutionary biologists because it represents an increase in complexity from individual to group-level reproduction and includes the evolution of a non-reproductive worker caste [10]. Along with reproductively-specialized castes, many of these societies have also evolved elaborate communication systems used to identify group members and to coordinate and divide labor among individuals [11]. These evolutionary innovations have afforded the social insects major ecological success – they are estimated to make up over 50% of the insect biomass on the planet even though they only account for ~2% of the insect species worldwide [12].

A great deal of effort has been focused on understanding the mechanisms that have enabled the multiple evolutionary origins of sociality [13]. Just like multiple cell types and tissues are derived from the same individual genome, the queen and worker castes are generated from a shared genomic background. This means that just as changes in gene expression drive cell type specifications, they should also drive developmentally-determined caste differentiation in the social insects. There is growing evidence to support this assertion, including well-documented differences in gene expression between castes in developing larvae and in adults [14–24], as well as differences in DNA methylation [25–28], post-translational histone modifications, and chromatin accessibility [29–31].

Although changes in coding sequences have been found to contribute to eusocial evolution in Hymenoptera [32], it is hypothesized that an expansion in the regulatory capacity of eusocial genomes may also have been a fundamental mechanism enabling these transitions [6]. This hypothesis is supported by a comparative study of 10 bee genomes that uncovered expansions in transcription factor binding sites in lineages where social behavior has evolved [6]. Similar observations have also been made in ants, both through examinations of non-coding sequence evolution [33] and by comparisons of gene expression patterns that have begun to uncover signatures of ancestral gene regulatory networks that may underlie caste determination [34]. However, direct comparisons of non-coding sequence evolution across species have not yet been leveraged to assess the contributions of these elements in the origins and elaborations of social behavior in bees.

Here, we identify non-coding regions that are alignable across eleven bee species that span three independent origins [35] and two independent elaborations of sociality [36] and over 100 million years of evolution. The alignability of these sequences across substantial evolutionary distances suggests that these regions are relatively conserved and that they could play a functional role in gene regulation. We have taken advantage of the convergent transitions in social behavior within bees to identify concordant evolutionary signatures in these non-coding sequences that are associated with the evolution of sociality. In general, we find that the landscape of these noncoding and putatively regulatory sequences in bees matches many of the patterns observed in conserved, non-coding elements (CNEs) in plants and vertebrates, including an exceptionally slow rate of evolution among those loci associated with genes involved in development. We then examine if and how this non-coding landscape has changed alongside behavioral and reproductive innovations associated with the evolution of eusociality. We find little association between non-coding sequence evolutionary rates and the origins of sociality across all bees, but we do identify several molecular pathways that have experienced convergent rate changes in association with the larger colonies and increased caste differentiation found within the stingless bees and honey bees. Finally, we discuss how these patterns of non-coding sequence evolution compare to patterns of coding sequence evolution and highlight future areas of research that can help to further illuminate the role of gene regulatory change in the evolution of eusociality.

## Methods

### Bee taxa included

We used previously published genomes for twelve bee species (Fig. 1; see Supplementary Information section 1.1 for detailed information on genome releases). For each species, we performed whole-genome alignments (see below) to identify non-coding alignable sequences (NCARs; excluding *E. dilemma*, for which the available genome sequence is highly fragmented). We also used this set of genomes to examine the evolution of protein coding sequence. Classifications of social behavior were drawn from previous studies [6], with respect to reproductive division of labor (SI 1.2). Species were split into four different behavioral categories: (1) solitary (*Dufourea novaeangliae, Habropoda laboriosa, Megachile rotundata*), (2) facultative simple sociality (*Ceratina calcarata, Eufriesea mexicana, Euglossa dilemma*, and *Lasioglossum albipes*), (3) obligate simple eusociality (*Bombus impatiens* and *Bombus terrestris*), and (4) obligate complex eusociality (*Apis florea, Apis mellifera*, and *Melipona quadrifasciata*; Fig. 1). Note that because of the variation in behaviors among species considered to have facultative simple social behavior, we refrain from using the more specific term “eusocial” to describe these species. Both obligate simple and obligate complex eusociality involve the presence of a queen and non-reproductive workers. However, the transition to complex eusociality, as designated here, involves an increase in the number of workers of at least several thousand, morphological specialization of castes, and vastly more complex systems of communication [37]. Simple sociality occurs in both obligately social and facultative forms wherein individuals vary in their expression of sociality within the species [38].

**Figure 1.**
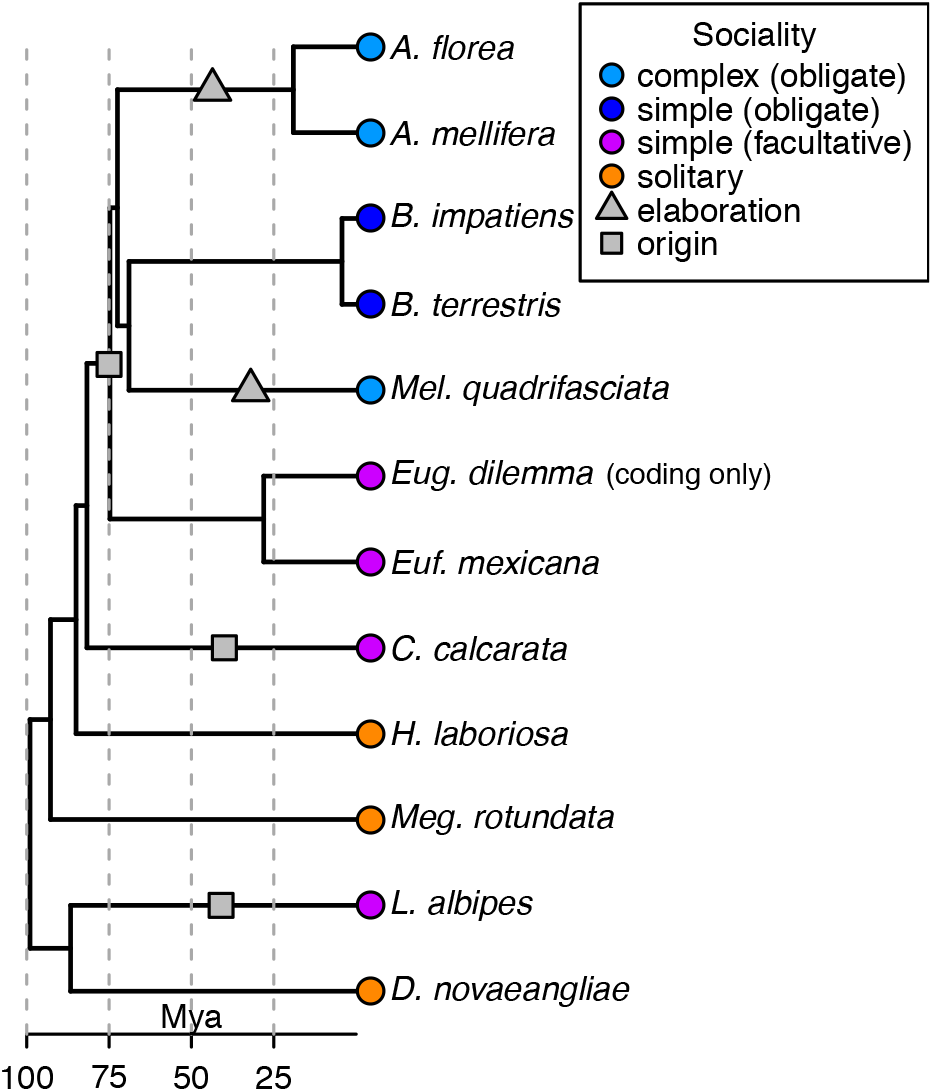
Phylogeny of bee species targeted in this study. These taxa span a range of behavioral forms, from solitary species that live and reproduce independently (orange), to eusocial species with a reproductive division of labor characterized by a queen and worker caste. Simple eusocial societies can either be facultative (purple) or obligate (dark blue). Complex eusocial (light blue) species contain nests made up of hundreds to thousands of individuals with morphological specializations between queen and worker castes. The species examined here encompass three independent origins of simple facultative eusociality and two independent origins of complex eusociality. Topology and dates are drawn from previous studies [36,82].

### Bee phylogeny

A large number of studies have explored the relationships among species in the family Apidae (represented by *Apis, Bombus, Eufriesea*, and *Melipona* in the current study). However, the most recent research shows that *Apis, Bombus*, and *Melipona* share a more recent common ancestor than any do with *Eufriesea* [36]. Thus, our study assumes that the ancestor of these three genera exhibited obligate simple eusociality and that there have been convergent elaborations of this behavior in the lineages leading to *Apis* and *Melipona*. Though it is possible that the ancestor possessed complex eusociality and that the lineage ancestral to *Bombus* reverted back to simple eusociality, because the transition from simple to complex eusociality is thought to be obligate and irreversible [11,39] and because no such reversals have been otherwise observed, such a scenario is less parsimonious.

### Identification of non-coding, alignable regions (NCARs)

Methods traditionally used for characterizing conserved, non-coding elements (CNEs) in vertebrates [40–43] rely on whole-genome alignments to assess changes in conservation in noncoding regions of the genome. However, the highly-fragmented nature of the publicly available bee genomes limits our ability to generate suitable whole-genome multiple sequence alignments that include all taxa in our study. To overcome this limitation, we instead relied on pairwise alignments of each species to the *Apis mellifera* genome to then generate multiple sequence alignments of non-coding regions as detailed below. Because these methods do not explicitly use sequence conservation, other than alignability, as a metric for identification, we refer to our sequences as non-coding alignable regions, or NCARs, rather than as CNEs. Given the relatively small fraction of the genomes (~0.5%; Table S1) of our target taxa that we were able to align, we concluded that these regions must be at least somewhat conserved compared with the rest of the genome sequence. Because we do not rely solely on extremely high levels of conservation across all species examined, our approach provides the benefit of allowing for the discovery of alignable non-coding regions whose rates of change are correlated with social evolution regardless of degree of conservation, thus allowing for the identification of potentially relevant regions that may not be identified using traditional CNE approaches. However, some NCARs may not be functional and/or subject to negative selection, potentially adding noise to our analyses.

Full genomes were first masked for repetitive sequences using RepeatMasker [44]. Each genome was then individually aligned against the *A. mellifera* genome using LAST [45,46]. The single best one-to-one alignment was used for each pair of sequences and non-syntenic regions were discarded (SI 1.3). Regions that were aligned against the same *A. mellifera* region across different species were merged together into a multiple sequence alignment and then realigned using FSA [47]. For our main analyses, we focused on only those regions for which at least nine species were represented. Coding sequences were masked for further analyses as we were focused on non-coding sequence. These resulting alignments were split into 500 base non-coding alignable regions (NCARs) for analyses, using a sliding window with 250 base step sizes so some NCARs overlap by 250 bases. Overlapping NCARs were excluded for the description of the distribution of NCARs across the genome and across feature types, discarding the NCAR with higher coordinates. Although our sliding window approach means that loci are not completely independent, it allows for fine-scale resolution of locations where rate changes have occurred. The resulting aligned sequence windows were filtered for quality using trimAl [48] to remove poorly aligned regions. Branch lengths were estimated for each taxon in each NCAR with BASEML [49,50] using the REV model of nucleotide substitution (model = 7) on a previously determined topology [36]. Full details of the alignment procedure are in SI 1.3 and scripts used to generate alignments are available at https://github.com/berrubin/BeeGenomeAligner.

### Functional classification and ortholog assignment

In order to assign putative functions to NCARs and to compare changes in the non-coding landscape to coding sequence evolution, we also identified single-copy orthologous genes in our dataset. These orthogroups were identified using ProteinOrtho v.5.15 [51] with a minimum connectivity of 0.5. Gene ontology terms were assigned to orthogroups using Trinotate [52] on *A. mellifera* representative sequences and gene names determined by orthology to *Drosophila melanogaster* genes found in OrthoDB [53]. When paralogous sequences from a species were detected in an orthogroup, all sequences from that species were discarded. Coding sequences were aligned using the coding sequence aware implementation of FSA [47]. Branch lengths for translations of all orthologous groups were then estimated with AAML [49,50] using the Empirical+F model of evolution (model=3). We estimated these branch lengths on the topology inferred previously [36].

We used HOMER [54] to identify *de novo* sequence motifs enriched in NCARs from each species, using the full genome of those species as background sequence. We then created a single set of motifs using those identified across all species using the compareMotifs.pl script. The resulting 147 motif seeds were used to identify similar motifs present in the NCARs for each species. When assigning putative function to motifs based on similarity to previously characterized binding motifs, we required a HOMER match score of 0.75 or greater. As transcription factor binding motifs have not been thoroughly characterized in any bees, these similarity matches are to motifs known from other, model organisms (e.g., *Drosophila*) and may not have the same functions in the taxa examined here.

We tested for differences in the abundance of motifs by comparing the proportions of NCARs that contained individual motifs in each species. We then compared these proportions between taxa with complex sociality and all other taxa using Wilcoxon rank-sum tests as well as Phylogenetic Generalized Least Squares (PGLS) tests assuming a Brownian motion error structure. To examine the presence of motifs across species within individual NCARs, we used χ^2^ tests, again comparing complex social taxa to all others. Although based only on the binary presence or absence of motif-detection, this approach is similar to those taken in previous studies in bees where motif matching scores were correlated with the evolution of social behavior [6]. Unfortunately, the small number of taxa included in these analyses and the large minimum p-values that result preclude the effectiveness of multiple test correction for either the tests for differences in overall motif abundance across taxa or the tests for differences in motif occurrence within individual NCARs. Although we believe that these results are useful as a starting point, they should be treated with caution because of these issues.

Putative functions were assigned to NCARs by the association with coding sequence in the genome of *A. mellifera* based on the midpoint coordinate of each NCAR. We split these gene-associated NCARs into sets associated with introns, 5’ UTRs, 3’ UTRs, promoters (<1.5kb upstream of the coding start site), and within 10,000 bases upstream or downstream of the coding start and stop coordinates. When NCARs were associated with multiple genes, they were assigned to individual genes based on the following priorities: 1. introns, 2. 3’UTRs, 3. 5’UTRs, 4. promoters, 5. upstream and downstream regions. When individual NCARs were present in the introns or UTRs of multiple genes, they were randomly assigned to a gene. When NCARs occurred in the promoters or upstream or downstream regions of multiple genes, they were assigned based on nearest proximity.

### Patterns of NCAR evolution

To assess the origins of novel non-coding elements in bees, we identified sets of NCARs that were present in all bee species, and those unique to five target clades including the obligately social corbiculates (*Apis, Bombus*, and *Melipona*), all corbiculates (the social corbiculates and *E. mexicana*), the corbiculates and *C. calcarata*, all Apidae (the corbiculates, *C. calcarata*, and *H. laboriosa*), and all Apidae and *Meg. rotundata*. For an NCAR to be considered unique to a clade, it had to be present in all species within that clade and have no recovered ortholog from any taxa outside of that clade. NCARs unique to clades were examined for possible functional enrichment by comparing the GO terms assigned to proximal genes and comparing these sets of GO terms with the set of NCARs used for the more general analyses (i.e. without specific requirements of taxonomy except that a minimum of nine species be represented). We also sought to compare the number of clade-specific NCARs across different clades while accounting for evolutionary divergence across taxa. We, therefore, inferred branch lengths for the overall phylogeny using a concatenated matrix of all protein sequences (SI 1.4). To standardize across clades, numbers of NCARs unique to each clade were multiplied by the total branch length inferred for that clade, thus downweighting closely related taxa and upweighting more distant relatives.

We also examined the overall rate of evolution in individual NCARs by standardizing the total branch length inferred for an NCAR locus to the branch lengths derived from the concatenated protein matrix while controlling for the taxa present in each NCAR. The resulting distribution of standardized non-coding branch length was examined to identify the fastest- and slowest-evolving NCARs across all taxa. The genes associated with these sets of NCARs were examined for GO term enrichment relative to the full set of NCARs present in at least nine species.

Previous studies in vertebrates have revealed that conserved, non-coding sequences tend to occur in genomic clusters [55]. To determine whether the same types of patterns are present in the NCARs of bees, we used permutation tests to identify significant clustering. These tests compared the number of 200kb windows with a minimum of 5, 6, 7, or 10 NCARs to the number of windows with a minimum of the given number of NCARs when locations of all NCARs were randomized within chromosomes (using 1,000 random permutations).

### Differences in evolutionary rates among bee species

We performed evolutionary rate tests on both NCARs and coding sequences to identify genomic regions that showed consistent changes in evolutionary rates associated with the evolution of social behavior using RERconverge [56–58] (SI 1.5). RERconverge calculates relative branch lengths by normalizing branches for a focal locus to the distribution of branch lengths across all loci. This enables analyses that look for convergent changes in evolutionary rates across different taxa while accounting for differences in phylogenetic divergence and in baseline rates of evolution across taxa. RERconverge compares rates of change in focal/foreground branches and the rest of the tree, and identifies loci that have a significant correlation between relative rates and a phenotype of interest. Slower rates of change among the focal branches can generally be interpreted as an increase in purifying selection among these taxa. Faster rates of change are more difficult to interpret as they may be indicative of either directional selection or a relaxation of purifying selection.

We made two different comparisons between social and solitary taxa. (1) We tested all taxa with any degree of reproductive division of labor against all other taxa (Fig. S1a). Note the inclusion of ancestral branches in these tests. (2) We identified NCARs and genes associated with the complex eusociality of *Apis* and *Melipona* by designating these terminal branches and the internal branch representing the ancestral *Apis* lineage as focal branches (Fig. S1b). The resulting sets of NCARs evolving significantly faster or slower on focal branches were examined for GO term enrichment among all genes proximal to NCARs represented by at least nine taxa using GO-TermFinder [59].

Although some previous work has examined molecular evolution in the obligately eusocial lineages (complex eusocial taxa + *Bombus*) we did not apply the relative rates test to this clade because the shared ancestry and single origin of eusociality is likely to generate a shared signal that would not be independent, reducing our confidence in the association between eusociality and the genes identified.

### Robustness of rate changes associated with social evolution

While RERconverge accounts for shared evolutionary history between taxa by treating each branch on the phylogeny as an independent data point, the currently available datasets create uneven sampling across clades that have evolved complex eusociality convergently (i.e., one *Melipona* lineage versus three *Apis* lineages including *A. mellifera, A. florea*, and the lineage ancestral to these two taxa). Thus, we were concerned that the majority of our signal was the result of lineage-specific evolution in *Apis*. To better assess this potential bias, we performed additional RERconverge tests using the three *Apis* lineages plus one of the two *Bombus* species as focal lineages instead of *Melipona*. Each *Bombus* species was tested separately. If *Apis* and *Melipona* share convergent rate changes related to complex eusociality, these loci should not show a significant association between *Apis* and either *Bombus* species which have simple eusocial behavior. *Bombus* is as closely related to *Apis* as *Melipona*, providing an ideal test case for characterizing the amount of signal contributed by *Melipona* to the RERconverge tests of complex eusocial taxa.

In addition, because current datasets include relatively few taxa, meaning that an outlying signal from a single taxon might have a drastic effect on our results (although the rank-based Kendall tests used by RERconverge partly remedies this issue), we used leave-one-out analyses to build confidence in our results, performing the RERconverge analyses using all iterations of two taxa among the three taxa with complex eusociality.

Next, we used a series of permutation tests to explore the degree to which our results were different from random expectations. First, we ran RERconverge on the full NCAR dataset using 1,000 sets of four randomly identified focal branches for assessing our test of complex eusocial lineages and using 1,000 sets of 13 randomly identified focal branches for assessing our test of lineages with any degree of sociality. P-value distributions resulting from our tests of complex eusocial lineages and all social lineages were compared to the distributions of p-values from these tests of random branches to determine if more loci were identified as significantly associated with social behavior more frequently than expected by chance. This approach for examining the enrichment of significant p-values is similar to that used previously for assessing the performance of RERconverge [56]. Results from these tests of random taxa were examined both for the numbers of NCARs evolving at significantly different rates and for GO term enrichment. We also generated null expectations for GO term enrichment by creating 1,000 sets of random NCARs equal in number to the number identified by RERconverge as significant in tests of complex eusocial taxa and all eusocial taxa. These NCARs were again tested for GO term enrichment.

Finally, we also explored the possible influence of gene tree discordance on our analyses of evolutionary rates but concluded that this phenomenon is unlikely to have substantially affected our results (SI 1.6, 2.11).

RERconverge, although shown to be a powerful method for detecting evolutionary rate changes associated with phenotype evolution [56,57], does not explicitly account for variation in GC-content, which has been found to influence evolutionary rate estimates in bees [6]. Thus, the results of the relative rates test may be influenced by variation in GC-content both within and between bee genomes. Future implementations of this type of test may benefit from the inclusion of GC-content as a factor, particularly among those taxa where this trait is known to vary widely across the genome, such as bees [60].

### Associations between NCARs and caste-biased gene expression

To investigate the relationship between non-coding regions and genes with caste-biased expression in *A. mellifera*, we drew lists of differentially expressed genes from three previous studies. We examined genes that were previously found to be expressed at different levels in virgin queens versus sterile workers for both adults [61] and larvae [62] as well as those differentially expressed between nurses and foragers, which represent categories of age-based worker polyethism [63]. Of the 3,610 genes compared between workers and queen adults, 587 were found to be worker-biased and 649 were found to be queen-biased by Grozinger et al. [61]. In a comparison of 4-day old larvae, He et al. [62] found that 276 of the 15,314 genes in the *A. mellifera* OGS v3.2 were worker-biased and 209 were queen-biased. Alaux et al. [63] compared 9,637 unique genes between foragers and nurses, 434 of which were expressed at greater levels in foragers and 464 of which were expressed at greater levels in nurses. Hypergeometric tests were used to test for enrichment of particular sets of genes.

## Results

### The landscape of alignable non-coding sequence in bees

Based on the results of our whole-genome alignments, we obtained 3,463 non-overlapping NCARs. Median divergence between *A. mellifera* and all other taxa across these NCARs varied from 4% in the most closely related *A. florea* to 17% in the most distantly related species (Table S1). Species representation across NCARs is given in Table S1. We used the genome of *A. mellifera* to examine the distribution of NCARs, finding that the vast majority (3,233) were present on scaffolds grouped into the 16 chromosomes of this species (Table S2) and were found in many regions associated with gene regulatory functions (Fig. 2a). In total, NCARs were within 10kb of 1,543 different genes. They were heavily enriched for proximity to coding sequence (hypergeometric test, p < 1×10^−20^), falling into one of the gene-associated categories 2.1-fold more often than expected by chance based on the proportion of the genome represented. 1,144 NCARs were in introns, 552 in downstream regions, 368 in promoters, 348 in 3’-UTRs, 249 in upstream regions, and 164 in 5’-UTRs (Fig. 2a). The remaining 638 NCARs were intergenic. 1,896 NCARs were within 10kb of multiple genes, 764 were within 1.5kb of multiple genes, and 88 NCARs overlapped UTRs or introns for multiple genes. Introns, UTRs, and promoters that contained NCARs tended to be longer and more GC-rich than all features of those types present in the *A. mellifera* genome (Wilcoxon rank-sum test, p < 0.01; Fig. S2). These characteristics are generally correlated with regulatory function [64,65], lending support to the hypothesis that NCARs may act as regulatory elements.

**Figure 2.**
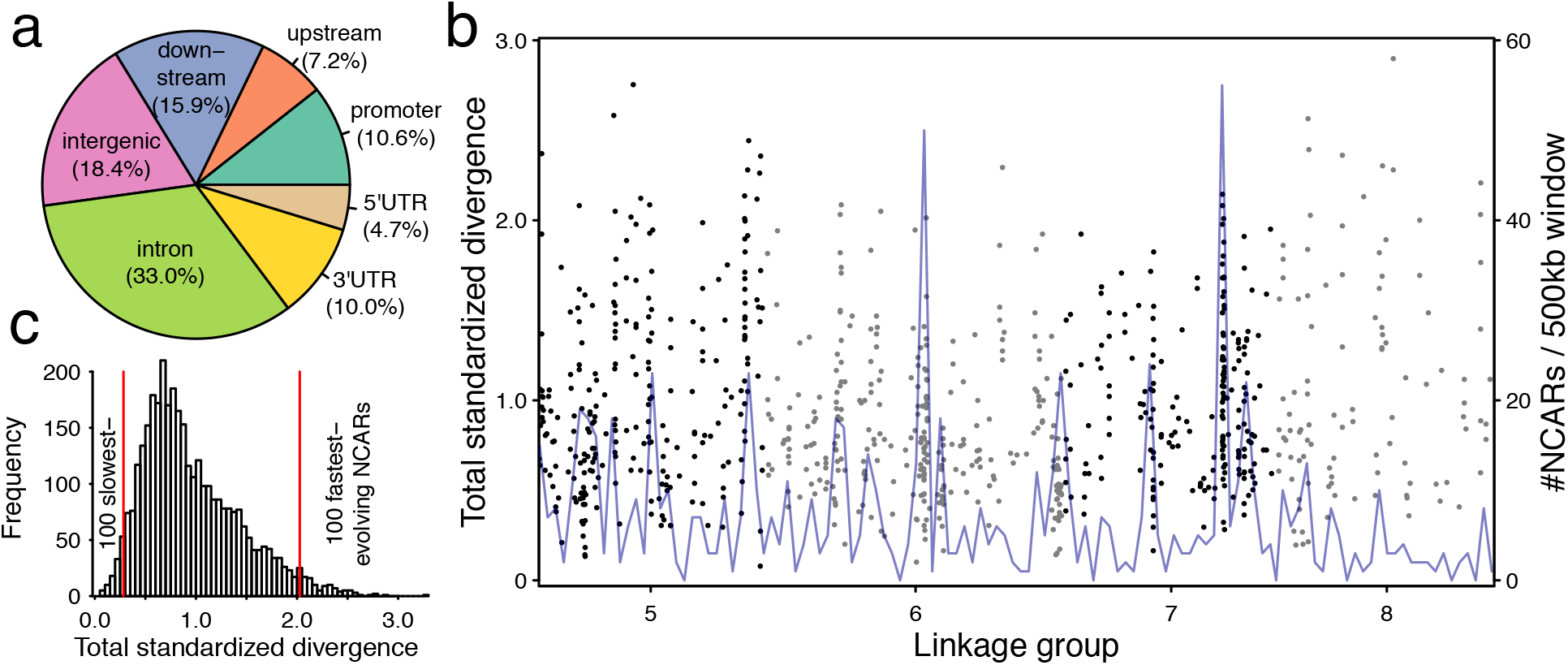
The landscape of non-coding alignable regions (NCARs) in bees. 3,463 NCARs were identified across bee genomes. Locations were mapped using the *A. mellifera* genome as a reference. (a) They were located in several genomic regions associated with gene regulation (see methods for classification scheme). (b) NCARs were distributed across all chromosomes, but are present in clusters on each chromosome. *A. mellifera* linkage groups 5-8 are represented on the x-axis; black and gray dots are used to denote each of these groups; each dot represents a single NCAR, and the y-axis signifies a standardized measure of divergence for each region (detailed in methods). The blue line denotes the # NCARs present in each 500kb window. NCARs occur in clusters across each chromosome, consistent with patterns observed in vertebrates and in plants. (c) NCARs exhibit substantial variation in evolutionary rates of change. The 100 slowest-evolving regions are associated primarily with regulation of gene expression, and the 100 fastest-evolving regions are associated with metabolism.

There were 532 NCARs present in all 11 bee genomes and, like many of the conserved noncoding elements in mammals [55], the genes proximal to these NCARs were enriched for GO terms related to developmental processes and transcription, relative to the full *A. mellifera* gene set (Table S3). Similarly, NCARs showed a significant clustering pattern across all chromosomes (permutation test p < 0.05; Figs. 2b, S3; Table S4), which is also typical of conserved non-coding elements in mammals [55] and plants [2]. All NCARs contained 56.6% AT on average, compared to the mean genomic background of 61.8% AT (Table S5). Rates of change in NCARs are significantly negatively correlated with GC-content (Pearson correlation p<1×10^−10^; Fig. S4).

Despite the different approach taken in our study, many of the characteristics apparent from studies of CNEs are also apparent among the NCARs identified here (clustering in the genome, association with developmental genes), suggesting that, although these results are not directly comparable, the two methods do identify related parts of the genome.

### The most rapidly evolving NCARs are functionally distinct from the most conserved

To examine general patterns of regulatory evolution across all bee species without considering differences in social behavior, we calculated the total standardized branch lengths (SI 1.5) for each NCAR and identified the top 100 fastest and slowest evolving regions (Fig. 2c). The 100 fastest evolving regions were associated with genes enriched for GO terms related to metabolic functions, while the 100 slowest evolving regions were associated with genes enriched for GO terms related to the regulation of gene expression (hypergeometric test, FDR-corrected p < 0.05; Table S6).

There were also differences in the types of genomic features associated with faster or slower evolving NCARs (SI 2.1). The fastest-evolving NCARs were enriched for presence in 5’ UTR sequence compared to the 3,233 non-overlapping gene-associated NCARs (hypergeometric test, p = 3.1×10^−10^, 4.5-fold enrichment), while the slowest-evolving NCARs were enriched in regions downstream of genes (hypergeometric test, p = 0.032, 1.64-fold enrichment). The fastest-evolving NCARs also contained 30% more binding motifs (n=3,763 occurrences of motifs proximal to 80 genes) than the slowest-evolving NCARs, which encompassed 2,973 motifs proximal to 69 genes. There were no major differences in which motifs were present in these sequences.

### Novel NCARs emerge alongside eusociality

Previous work in vertebrates has suggested that the origin of novel phenotypes is correlated with the appearance of novel clusters of CNEs associated with distinct types of genes. For example, before mammals split from reptiles and birds, CNEs were recruited near transcription factors and their developmental targets, but CNEs that arose in placental mammals are enriched near genes that play roles in post-translational modification and intracellular signaling [40].

To determine if similar recruitment processes have played a role in the evolution of eusociality, we identified NCARs that are unique among the social corbiculates (*Apis, Bombus, Melipona*, and *Eufriesea*). The recruitment of novel NCARs in this group may indeed be associated with their shared origin of sociality. We found 1,476 NCARs associated with 605 genes that are shared among all of these species and unique to this clade (Fig. 3a). Although neutral expectations would predict that the clade containing only *Apis, Bombus*, and *Melipona* would contain the greatest number of NCARs, the clade including *all* corbiculates contained the largest number of clade-specific NCARs (both raw and standardized by total clade branch lengths, and despite the fact that *E. mexicana* is one of the most fragmented genomes in the dataset [6]), suggesting that there was an expansion in regulatory regions at the origin of this clade. Genes proximal to these regions are not enriched in particular functions after multiple-test correction, although many nervous system functions show some indication of enrichment (hypergeometric test, uncorrected p < 0.01; Table S7). These corbiculate-specific NCARs are located primarily in introns (35%) and intergenic regions (21%), similar to the distribution of all NCARs.

**Figure 3.**
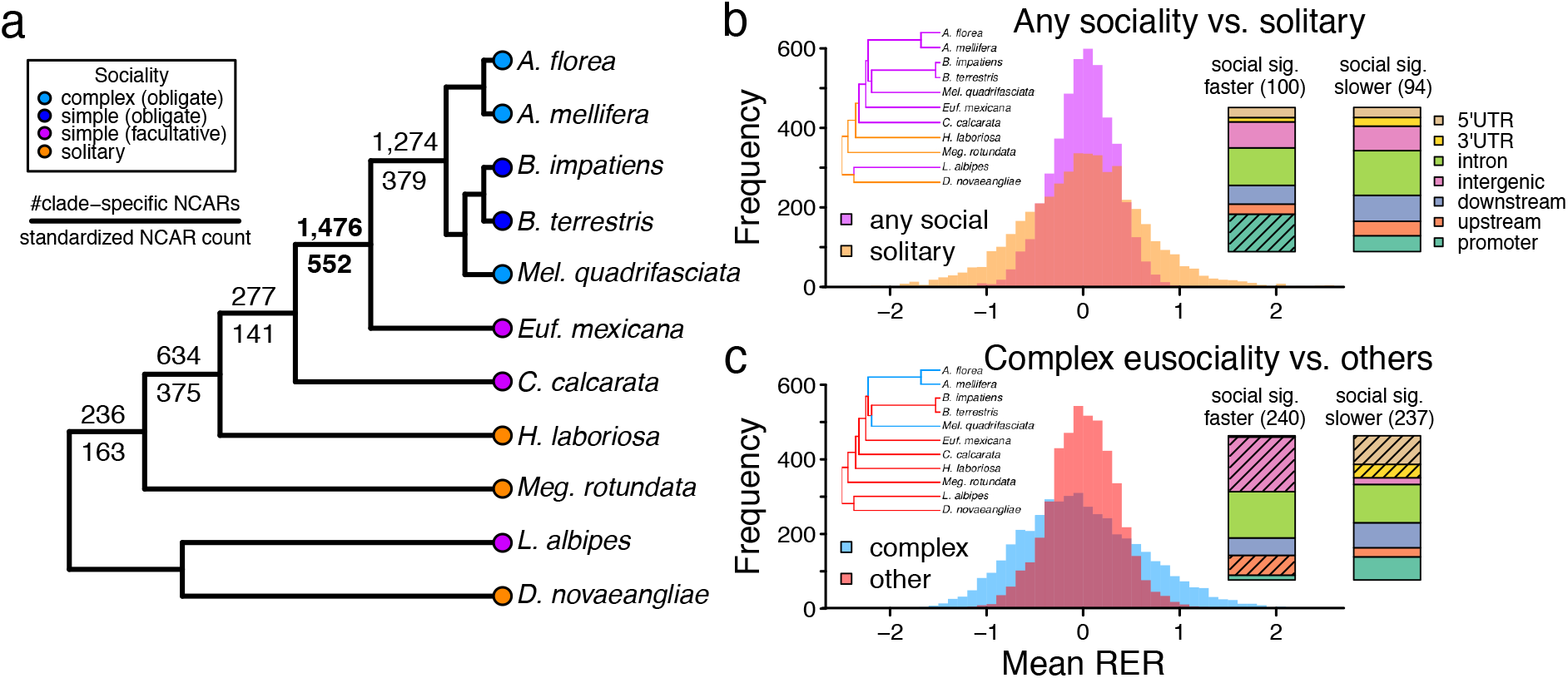
NCAR evolution correlates with evolution of eusociality. (a) Novel NCAR recruitment is associated with the emergence of obligate eusociality in the corbiculate bees (bolded text). 1,476 NCARs are shared uniquely among these species and are enriched for gene functions associated with cell and nervous system development. NCAR counts below branches are standardized by multiplying by total branch length within the clade. (b) Distribution of mean relative rates among taxa with any degree of sociality vs. strictly solitary taxa in all NCARs. 171 NCARs show signatures of convergent evolution across all social bee species relative to solitary taxa. 100 of these are evolving more rapidly while 94 are changing more slowly. Fast-evolving regions are enriched for promoter sequences (inset; hypergeometric test, p=4.5×10^−5^), and contain a surplus of *Fmr1* binding motifs. (c) Similarly, distribution of mean relative rates among taxa with complex sociality vs. others in all NCARs. Branches treated as foreground and background are shown in the inset phylogeny. There are 477 NCARs that show convergent rate changes on complex eusocial branches. Rapidly evolving regions are associated with neuronal fate, and are located in upstream and intergenic regions more often than predicted by chance (hypergeometric test, p<1.0×10^−5^). Shading indicates significantly enriched feature types.

### A subset of NCARs show concordant rates of change associated with sociality

*Convergence across bee species that exhibit any form of reproductive division of labor*. Our dataset encompasses three independent origins of reproductive division of labor (sociality) in bees (Fig. 1). To be sure that potentially important functional regions were not divided across NCAR loci, we included the full set of 4,611 NCARs in our analysis, including NCARs that overlapped in sequence. Of these, 4,582 loci had the requisite taxon composition to be included in the relative rates test. We found 100 NCARs with signatures of accelerated evolution in all social relative to non-social bees and 94 with deceleration (relative rates test, p < 0.05; Table S8). The distributions of mean relative rates for all social and all solitary taxa across all NCARs were similar, showing that our tests were not biased to find significance in a particular direction (Fig. 3b). Note that the difference in variance between distributions is most likely due to the larger number of social than solitary taxa and may increase the chances of spuriously identifying significant rate changes.

The number of loci with significant rate changes associated with all social lineages (p < 0.05; 4.3%) is not more than would be expected by chance; the p-value distribution of all loci from these tests is similar to that resulting from tests of 1,000 permutations of randomly selected lineages (Fig. S5a). These permutations yielded at least the same number of significant loci 565 times (56.5%). Consistent with this pattern, the genes proximal to the NCARs evolving at different rates in social species were not significantly enriched for particular functional gene classes after multiple test correction (Table S9), although NCARs evolving faster in social taxa were found more frequently in promoters than expected by chance (when compared with the set of NCARs included in the relative rates test; hypergeometric test, p=4.5×10^−5^, 2.4-fold enrichment; Table S8; Fig. 3b inset). No association with gene features was found for the NCARs evolving at a slower rate in social taxa. In general, promoters are thought to experience greater levels of evolutionary constraint relative to other regulatory features, and this higher degree of conservation may help to explain why we can identify larger numbers of loci with concordant signatures of selection across the largest evolutionary divergences in these regions [66].

*Bee species representing independent origins of complex eusociality*. The honey bees (*Apis*) and the stingless bees (*Melipona*) share a eusocial ancestor, but most likely represent two, independent transitions from simple to complex eusociality (i.e. with morphologically specialized castes, swarm-founding, and large colony sizes) within the social corbiculates [6,67]. We tested these complex eusocial lineages for significant differences in evolutionary rates compared to all other taxa. Again, distributions of mean relative rates were not skewed by behavioral type, so our results should not be biased to identify changes in evolutionary rates in one particular direction (Fig. 3c) and differences in variance are likely due to differences in sample size across behavioral groups. In contrast to the above tests encompassing species with any form of division of labor, the distribution of p-values obtained from tests for an association with complex eusociality were enriched for low values (11% had p < 0.05) relative to the p-values obtained from tests of 1,000 random sets of branches (Fig. S5b). 4,287 NCARs had the required taxon composition for inclusion in the test, 240 of which exhibited faster rates of evolution in these complex eusocial lineages relative to all other bees (relative rates test, p < 0.05; Table S8). These were associated with genes enriched for a total of nine GO terms, including neuron fate and differentiation (hypergeometric test, FDR-corrected p < 0.05; Table S9, S10) and were found more often than expected by chance in upstream and intergenic regions compared to the set of all NCARs included in the relative rates test (hypergeometric test, p < 0.01; Fig. 3c; Table S8). Similarly, there were 237 NCARs evolving at significantly slower rates in complex eusocial taxa compared to all other bees (relative rates test, p < 0.05; Table S8). The genes proximal to these NCARs were not significantly enriched for any GO terms after multiple test correction (hypergeometric test, FDR-corrected p > 0.05; Table S9). These NCARs were found more often than chance in 5’ and 3’ UTRs (hypergeometric test, p < 0.01; Table S8).

To determine whether these results are robust to taxon sampling, we ran the relative rates test on subsets of complex eusocial taxa. While the results are, as expected, much weaker, eight of the nine GO terms enriched in the test of all complex eusocial lineages also show signatures of enrichment in at least one of these tests of subsets of taxa (uncorrected p < 0.05; SI 2.2, Table S11). We also identified fewer loci with convergent signatures of rate changes between *Apis* and *Bombus* lineages (369 in *B. impatiens* and 360 in *B. terrestris* versus 473 in the test of *Apis* and *Melipona*) confirming that a greater number of NCARs evolve in parallel across complex eusocial lineages than between these complex and simple eusocial lineages (SI 2.3). None of the nine GO terms enriched in the test of all complex eusocial lineages are significantly enriched in tests combining *Apis* and either *Bombus* lineage (hypergeometric test, FDR-corrected p > 0.3).

The 1,000 permutations of RERconverge using four random foreground lineages also supported the results from our test of complex eusocial taxa, showing that our results differed from random expectations. These tests based on random foreground lineages had medians of 99 NCARs evolving significantly faster and 99 NCARs evolving significantly slower. The 99^th^ percentiles were 178 and 160 for faster and slower evolving NCARs, respectively. None of the 1,000 permutations yielded at least 240 faster evolving loci, the number of significantly faster evolving loci resulting from the test of complex eusocial lineages. Only a single permutation yielded at least 237 slower evolving loci, the number of significantly slower evolving loci from the test of complex eusocial lineages. Thus, the test for loci evolving at different rates in complex eusocial taxa finds significantly more loci with rate changes than expected by chance (permutation test, p ≤ 0.001).

We also examined the sets of significantly faster evolving NCARs in each of these random permutations for GO term enrichment and found an average of only 0.06% of GO terms tested were significantly enriched versus 1.0% in the 240 NCARs identified as evolving significantly faster in complex eusocial taxa. Thus, the random expectation is that 0.5 GO terms will be identified as significantly enriched by chance whereas nine terms were identified in tests of complex eusocial lineages, suggesting a strong, non-random association. In addition, these nine GO terms were identified as significantly overrepresented no more than 3 times among the 1,000 permutations of random lineages, showing that each of these nine terms is rarely identified by chance (permutation test p ≤ 0.003; SI 2.4). Thus, multiple approaches demonstrated that our tests for convergent evolution among taxa with complex eusociality yielded results that differed from random expectations, providing confidence in our results and analytical framework.

### Sequence motifs associated with social evolution

NCARs showing concordant rate changes across all forms of social behavior contained a similar number of known motif occurrences regardless of whether these regions were faster or slower evolving in social relative to solitary lineages (n=3,235 in faster NCARs and 2,842 in slower NCARs). Ignoring any signature of evolutionary rate changes, there were four motifs that were significantly more abundant in the NCAR sequences of social bee taxa relative to other branches (PGLS p < 0.05; SI 2.6; Table S12). One of these was a *Drosophila* binding motif for the Fragile X protein gene, *Fmr1*, a gene known to play a key role in brain development across a wide range of animals [68] and previously associated with social evolution in bees [6].

As we found with all social lineages, NCARs associated with the evolution of complex eusociality contained a similar number of sequence motif occurrences regardless of whether they were fast or slow evolving (n=9,677 for faster regions and 9,311 for slower regions; SI 2.5). Thus, there is not likely to be a simple increase in the number of motifs present in accelerated regions relative to those that show increased constraint. We also found little evidence for changes in motif abundance in those NCARs associated with social evolution (SI 2.6; Tables S13, S14).

### NCARs are not associated with gene expression differences among castes

Genes differentially expressed between honey bee castes are not generally overrepresented in NCAR-associated genes (hypergeometric test, p > 0.05; SI 2.9). However, there were 15 NCARs proximal to 11 different genes that showed convergent acceleration associated with the elaborations of eusociality in honey bees (*Apis*) and stingless bees (*Melipona*) that were previously shown to be differentially expressed between castes in honey bees (Table S15). Similarly, there were 21 NCARs with slower rates of evolution on the branches associated with the elaboration of eusociality that have also been shown to be differentially expressed in socially-relevant phenotypes in honey bees.

### Both NCARs and coding-sequences show signatures of convergent evolution, but on different functions

The same relative rates tests used to identify changes in NCARs can also be used to identify changes in coding sequence, and we uncovered 10 genes that showed concordant increases in rates on all social branches and on all complex eusocial branches. There is a significant overlap in both genes and GO terms between our study and a previous study [6] that used different methods to identify signatures of selection across this group of bees (calculated based on the number of overlapping genes showing concordant changes on complex eusocial branches; hypergeometric test, p = 0.0004, 2.0-fold enrichment; SI 2.10).

Overall, we find that coding sequence and NCAR sequence evolution appear to be quite distinct. We find no correlation between total standardized branch lengths between NCARs and proximal genes, regardless of the distance of NCARs to genes (log_2_-transformed R = 0.04 p = 0.50; Fig. S7), as well as when limited to just introns (log_2_-transformed R = −0.03, p = 0.81) or 3’ UTRs (log_2_-transformed R = 0.12, p = 0.33). Moreover, the genes and functional terms associated with changes in NCAR rates are distinct from the genes and functional terms associated with evolutionary changes in coding sequence. For example, although NCARs evolving more slowly in complex eusocial taxa show no GO term enrichment (Table S9), slowly-evolving protein-coding sequences are enriched for small molecule transport and catabolism (Table S16). And protein-coding genes evolving more rapidly in complex eusocial lineages are associated with cell projections (Table S16), while NCARs evolving more rapidly in complex eusocial lineages are associated with cell fate commitment and neuron differentiation (Table S9). Although processes associated with cell projections among the protein-coding genes may include or overlap with neuronal development, NCARs are clearly enriched in this type of process to a greater degree. This suggests that the changing selective pressures that occur during the evolution of eusociality may act on the regulatory elements and protein sequences of different sets of genes.

However, of the 317 genes included in both the NCAR and coding sequence tests of rate differences in complex eusocial lineages, there were 6 genes that showed consistently slower rates of change in both (hypergeometric test, p = 0.0049, 3.3-fold enrichment; Table S17) and three genes that showed consistently faster rates of change in both (hypergeometric test, p = 0.046, 3.5-fold enrichment; Table S17). This overrepresentation indicates that rates of evolution are concordant between some coding and proximal non-coding sequences, although this may only occur when loci are subject to stronger selective pressures.

### No apparent bias of selection on regulatory versus coding sequence

It is possible that the origins of sociality are associated primarily with changes in gene regulation rather than with changes in coding sequence evolution [69]. However, we did not find any evidence that the proportion of NCARs with evolutionary rate changes associated with sociality was greater than that found in coding sequences (Table S18). As expected from relative rates inferences, we did not find any apparent differences in the distributions of evolutionary rates in the focal or background lineages of coding and non-coding sequences (Figs. 3, S8). However, the total standardized divergence (total branch lengths for a locus standardized by number of taxa and nucleotides) was greater in NCARs than in CDS, as expected when comparing non-coding to coding sequence evolution (Wilcoxon rank sum test, p < 1×10^−10^; Fig. S9). That said, non-coding and coding sequences do overlap in their distributions (Fig. S9), demonstrating that in bees, as in other taxa [70], some non-coding sequences can experience the same level of constraint as protein-coding sequences.

## Discussion

### The landscape of putative regulatory sequences in bees is similar to mammals and plants

We have characterized a landscape of putatively regulatory non-coding sequences in bees. Consistent with the theory that these non-coding landscapes may have ancient, metazoan origins [1], we have found that the features of this landscape are similar to those described in vertebrates [55] and plants [2]. We find that NCARs are distributed throughout the genome in clusters, and those regions that are present in all bee species examined are enriched for developmental functions.

### Regulatory innovations are associated with the evolution of eusociality

Many of the major evolutionary innovations in vertebrates have been linked to the appearance of novel clusters of conserved non-coding elements [40], and each innovation appears to be associated with different types of gene functions. We initially predicted that the greatest gain in NCAR number would have occurred in the ancestor of the obligately eusocial clade (*Apis, Bombus*, and *Melipona*), in part because our use of *A. mellifera* as a reference for genome alignments was expected to bias NCAR discovery towards the closest relatives of this species. However, even after standardizing NCAR counts for evolutionary divergence time, the more expansive clade of corbiculate bees (*Apis, Bombus, Melipona* and *Eufriesea*), which share a simple eusocial ancestor, has the largest number of clade-specific NCARs. These results suggest that the origin of eusociality in this clade was accompanied by an increased regulatory capacity provided by these NCARs.

### There are concordant changes in non-coding sequences associated with sociality

Although the regions that show concordant rate shifts on all social lineages may represent changes that are important in the establishment of sociality, several lines of evidence presented above suggest that many of the significant changes detected are likely spurious. However, the NCARs associated with the elaborations of eusociality in honey bees (*Apis*) and stingless bees (*Melipona*) appear to represent a true signal of convergent rate changes. Faster evolving sequences on these branches were enriched for sequences upstream of genes and were associated with genes that play important roles in neuron fate commitment as well as a number of developmental processes. Loci with rate shifts in complex eusocial taxa include at least two NCARs located within introns of genes (the intron of *Fmr1* has slower rates and the intron of *ftz-f1* has faster rates) previously associated with social behavior and known to play key roles in neuronal remodeling and development of the mushroom bodies [71,72] (Fig. S6). This is a brain region crucial for sensory integration and learning and memory in insects, and is thought to play an important role in caste differentiation in honey bees [73,74]. Higher rates of change in complex eusocial taxa in *ftz-f1* and other loci likely indicate either a loss of function and concordant relaxation in purifying selection, directional selection acting to change the regulatory activity of the region, or some combination of the two: previous regulatory action may be eliminated while selection simultaneously acts to construct new binding sites or functions, changing the way the associated genes are expressed. Lower rates of change as seen in the intron of *Fmr1* may instead indicate increased purifying selection and a maintenance of consistent function. Regardless, changes in these non-coding sequences may influence neurodevelopmental and other processes and, thereby, the evolution of social behavior.

In addition, we were able to identify binding motifs present at significantly higher frequencies in regions evolving more rapidly in complex eusocial taxa, as well as motifs that occurred at higher frequencies in regions evolving more slowly in complex eusocial taxa relative to all other species. As with the above results examining the origins of sociality in bees, these results also provide evidence that similar transcription factors or binding proteins may have been co-opted by both honey bees and stingless bees as eusociality increased in complexity in each of these groups.

### Little evidence for an association between NCAR evolution and caste-biased gene expression

Because at least some of the characterized NCARs are likely to represent functional regulatory elements, we predicted that these regions might be enriched for proximity to genes whose expression has previously been associated with caste differences in social lineages. Indeed, we did identify some NCARs whose evolutionary rates were associated with sociality that were also proximal to a number of genes known to exhibit expression differences among honey bee castes (e.g., *Fmr1* [68], *Sema-1a* [75,76], *babo* [77,78], *ftz-f1* [71,79], and *shep* [80]; Table S15). However, we failed to find a significant overall enrichment of NCARs proximal to caste-biased genes.

A number of methodological issues may influence this finding. First, only a small subset of the tested differentially expressed (DE) genes in honey bees were also associated with NCARs and included in our dataset, making it difficult to generate a robust statistical inference. While this could represent a true lack of overlap, it could also be an artifact of the EST-based microarrays that several of these DE sets used, and coupled with the approaches we implemented to identify NCARs, we may be missing substantial proportions of genes that would show these concordant signatures. Alternatively, because the gene expression datasets available are primarily limited to honey bees while the comparisons we are making are across multiple species, many of the genes we identify may not have as large-scale expression differences as those that are species-specific. Both novel and conserved genes are differentially expressed among eusocial insect castes [22], yet our approach would only conceivably identify NCARs proximal to those which are at least somewhat conserved. Finally, within the honey bees, most large studies compare differences between adult bees [61,63], while the NCARs we have identified could affect gene expression at any point throughout development, and it is difficult to predict when, where, and in what context gene expression changes may occur. Although we did examine overlap with genes differentially expressed between worker and queen larvae (SI 2.9), these results were based on a relatively small dataset and may have only captured those genes with the most extreme expression differences [62]. Additional large-scale studies of expression differences across developmental stages and specific tissues will be necessary to draw strong conclusions on the association between NCARs and genes fundamental to social behavior.

### Evolutionary dynamics of non-coding and protein-coding sequences

We have used the same statistical analyses to examine and compare both coding sequence and NCAR sequence evolution. In general, we find no evidence to support the idea that a greater proportion of NCARs than coding sequences have experienced novel selective pressures associated with the evolution of sociality. It should be noted that our analyses focus on concordant evolutionary signatures in regions that are alignable across species. As a result, our dataset and analyses cannot examine the role that novel regulatory regions (i.e., regions that are unique to individual taxa) may play in the evolution of sociality. This kind of regulatory innovation could indeed be a key feature associated with the origins of sociality, but is beyond the ability of our current datasets and analyses to detect. We did observe an increased number of alignable, noncoding sequences associated with the origin of eusociality in the corbiculates, providing a glimpse into the potential role that regulatory novelty may play in this process. However, future work is needed to better characterize novel regulatory elements, many of which are likely to be taxon-specific.

Remarkably, some NCARs are evolving at the same overall rate as the most conserved coding sequences, suggesting that, at least for some of the non-coding regions that we can align across species, negative selection may be just as strong as it is for some proteins. Although our results are not directly comparable, they echo the results of mammalian studies, where non-coding, ultraconserved elements (UCEs) show similar or stronger levels of negative selection than many coding sequences [70].

### Limitations of this study

This study has focused on a small subset of bee species for which genomic resources have already been developed. These species are heavily biased towards social lineages, and thus most of the comparative power comes from the corbiculate bees, which share a single origin of sociality. Moreover, these taxa span large periods of evolutionary divergence, and the analyses we have implemented here have been based primarily on sequence conservation among these different taxa. There are over 20,000 bee species on this planet, and there have been up to 5 independent origins of sociality within this clade [81]. Future work focused on more closely-related lineages that encompass more of these evolutionary transitions can help provide greater insight into the role of gene regulation in the origins of sociality.

A number of technical limitations also limit the power and completeness of our study. Most glaring is the high variability in quality of the genome sequences used. Because of these limitations, we have focused on alignable non-coding regions rather than those that are especially highly conserved (as has been done previously [1,3,33,40]). Although this approach enables the examination of a broader palette of sequences, it also creates several difficulties. For example, our approach will fail to detect regulatory sequences that are both not sufficiently conserved as well as those that do not appear in a sufficient number of genome sequences as a result of incomplete assembly. Thus, we almost certainly failed to detect large numbers of alignable sequences simply due to the draft nature of the genomes included. Moreover, the identified NCARs are not necessarily functional or subject to negative selection, nor are neighboring NCARs statistically-independent, and it is possible that non-homologous sequences could be included in some cases. All of these factors contribute to background noise in the analyses we have presented and reduce our ability to detect loci evolving in association with social behavior.

Despite these limitations, our methods have succeeded in identifying several promising associations between non-coding sequences and social evolution in bees. We hope that this work can help to spotlight the benefits of research into non-coding sequence evolution and to motivate the generation of additional genomic resources for social insects and similar model systems.

## Conclusions

Changes in non-coding sequences are likely to play an important role in the evolution of sociality. We find that a large number of non-coding regions have been recruited alongside the origin of simple eusociality in corbiculate bees, highlighting a possible role in this behavior. Moreover, we observe concordant changes in alignable non-coding sequences associated with two transitions from simple to complex eusociality. Thus, the analyses of non-coding regions in this study have helped to uncover convergent signatures of social evolution that would have otherwise been overlooked by investigation of coding sequence alone. These results highlight the utility and importance of examining both coding and non-coding change to understand the molecular mechanisms underlying phenotypic evolution.

## Acknowledgements

We thank Micah Fletcher, David Galbraith, Luke Henry, Luisa Pallares, Lance Parsons, Serge Picard, and Eli Wyman for their feedback and support. Nathan Clark, Wynn Meyer, and Raghavendran Partha provided valuable guidance for using the relative rates test.

## Data accessibility

NCAR sequences and genomic coordinates and the main analytical pipeline are available from GitHub: https://github.com/berrubin/BeeGenomeAligner.

## Authors’ contributions

BERR, BGH, and SDK conceived the project. BERR performed computational analyses. BMJ compiled gene expression datasets. BERR and SDK drafted the manuscript, and all authors revised and approved the final version.

## Supplementary figures

**Figure S1.** Phylogenies used to conduct relative rates tests with focal lineages colored in red.

**Figure S2.** Distributions of GC-content (top row) and lengths (bottom row) of sequence features in which NCARs were identified (red) and all sequence features (blue) in the *A. mellifera* genome. P-values are the result of Wilcoxon rank-sum tests comparing these distributions. The length distribution of promoters is not shown because promoter length was fixed at 1.5kb.

**Figure S3.** NCAR distribution across *A. mellifera* linkage groups 1-4 and 9-16 are represented as in Fig. 2b. Dots show the locations of NCARs. Black and gray colors are used to denote the linkage groups and the y-axis signifies a standardized measure of divergence for each region (detailed in methods). The blue line denotes the # NCARs present in each 500kb window.

**Figure S4.** GC-content of *A. mellifera* sequence in each NCAR as a function of standardized total branch length of all taxa present in the NCAR.

**Figure S5.** Distribution of p-values obtained from relative rates test including all lineages with any degree of sociality as focal taxa (a) and from relative rates test focused on only those lineages with complex eusocial behavior (b). Red bars show the results from the test of the indicated focal lineages and blue bars show the p-values obtained from 1,000 iterations of relative rates tests on randomly chosen focal lineages.

**Figure S6.** Two intronic NCARs associated with complex social behavior are key regulators of mushroom body neuronal remodeling (*ftz-f1*; [71]) and development (*Fmr1*; [72]). *ftz-f1* shows accelerated rates of change on complex social branches relative to the remaining branches in the tree (relative rates test, p=0.008). *Fmr1* shows significantly slower evolution on complex social branches (relative rates test, p=0.009).

**Figure S7.** Log-transformed total branch length of coding sequences and proximal NCARs standardized to the branch lengths inferred from all a concatenation of all protein sequences. When multiple NCARs were associated with individual genes, mean standardized branch lengths were used.

**Figure S8.** (a) Distribution of mean relative rates among taxa with complex sociality vs. others in all coding sequences. (b) Distribution of mean relative rates among taxa with any degree of sociality vs. strictly solitary taxa in all coding sequences.

**Figure S9.** Distribution of total evolutionary change in all CDS’s and NCARs analyzed. To make these measures comparable across loci and sequence classes, the standardized total evolutionary change was additionally divided by the number of bases in each locus.

## Supplementary tables

**Table S1.** Species representation in 3,463 non-overlapping NCARs.

**Table S2.** Distribution of NCARs across the 16 *Apis mellifera* chromosomes.

**Table S3.** GO terms enriched in genes proximal to NCARs present in all 11 bee taxa.

**Table S4.** Permutation tests of NCAR clustering in 200kb windows.

**Table S5.** Mean AT-content of NCARs.

**Table S6.** GO terms enriched in genes proximal to the 100 fastest- and slowest-evolving NCARs.

**Table S7.** GO enrichment in clade-specific NCARs.

**Table S8.** Sequence features of NCARs identified as associated with the evolution of sociality.

**Table S9.** GO enrichment in genes proximal to NCARs associated with sociality using RER tests.

**Table S10.** Genes involved in neuron differentiation proximal NCARs evolving faster in taxa with complex sociality.

**Table S11.** Enrichment of the nine GO terms identified as significantly enriched in NCARs evolving significantly faster in complex eusocial taxa when individual taxa were excluded from analyses.

**Table S12.** Sequence motifs that differ in abundance in species with any degree of sociality.

**Table S13.** Sequence motifs that differ in abundance in species with complex sociality.

**Table S14.** Motifs that differ in frequency in NCARs associated with complex social taxa by RER test.

**Table S15.** Genes with both caste-biased expression and proximal NCARs with exceptional rates of evolution.

**Table S16.** GO enrichment in genes associated with sociality using RER tests.

**Table S17.** Genes with significantly different rates of evolution in both coding and proximal non-coding sequence.

**Table S18.** Numbers of coding and non-coding sequences evolving at significantly different rates.

**Table S19.** Motif abundances across all taxa and results of Wilcoxon tests comparing abundances between complex eusocial taxa and all other taxa.

