## Supplementary Information for "Rate variation in the evolution of non-coding DNA associated with social evolution in bees"

### 1. SUPPLEMENTARY METHODS

#### *1.1. Genome sequences*

We examined 11 previously published genomes (Fig. 1) in order to identify conserved non-coding sequences. These included *Apis mellifera* (genome version 4.5, OGS v. 3.2)*, Apis florea* (genome version 1.0, NCBI annotation release 101)*, Bombus terrestris* (genome v. 1.0, OGS v. 1.3)*, Bombus impatiens* (genome v. 2.0, OGS v. 1.2)*, Ceratina calcarata* (NCBI annotation release 100), *Dufourea novaeangliae* (OGS v. 1.1)*, Eufriesea mexicana* (genome v. 1.0, OGS v. 1.1)*, Habropoda laboriosa* (genome v. 1.0, OGS v. 1.2)*, Lasioglossum albipes* (genome v. 2, OGS v. 5.42)*, Megachile rotundata* (OGS v. 1.1)*,* and *Melipona quadrifasciata* (genome v. 1.0, OGS v. 1.1) [1–6]. No versions have been provided for the genome assemblies of *D. novaeangliae* or *M. rotundata.* We also explored coding sequence evolution in all of these species plus *Euglossa dilemma* [1]. Data for all species was downloaded from BeeBase except for *A. florea* which was downloaded from GenBank (GCF_000184785.2_Aflo_1.0)*, C. calcarata* which was also obtained from GenBank (GCF_001652005.1_ASM165200v1), and *E. dilemma* which was downloaded from http://www.eve.ucdavis.edu/sanram/scripts.html. For *A. florea* and *C. calcarata*, the RefSeq annotations were used for coding sequence evolution analysis. We drew *A. mellifera* miRNA annotations from miRBase v22 [7]. We did not include *E. dilemma* in analyses of non-coding sequences because the highly fragmented nature of this genome made alignments difficult and potentially unreliable.

Although all genomes and annotations included in this study are relatively complete based on analyses of coding sequences [3], the annotations for *A. mellifera* are likely of greater quality than for other taxa, as this species has acted as a genomics model for more than a decade. Harder to assess than completeness of coding sequences is the completeness of non-coding sequences and we expect that these regions are also likely better-represented in the *A. mellifera* genome. While we do not predict that these differences in genome quality will produce any serious biases in our analyses, the availability of a greater number of very high quality genomes would clearly be beneficial, allowing us to more accurately identify larger numbers of non-coding alignable regions across taxa.

##### *1.2 Classification of eusociality*

We used Kapheim et al. [3] to classify taxa into different types of sociality. These designations are based on a number of characteristics including colony size, degree of morphological difference between reproductives and workers, and the presence of social polymorphism [8,9]. In our dataset, we consider the two honey bee species, *Apis mellifera* and *A. florea*, as well as the stingless bee species, *Melipona quadrifasciata*, to be representative of "complex" eusociality. These taxa have distinct worker and reproductive castes, have colony sizes reaching tens of thousands of individuals, and show no social polymorphism (i.e., all individuals of these species are social). The two *Bombus* species, *B. impatiens* and *B. terrestris*, are also obligately social but colony sizes reach only into the hundreds and there is little morphological differentiation between workers and reproductives, so we consider these species to exhibit "simple" sociality. *Euglossa dilemma* exhibits a form of simple eusociality but this trait varies across the species [1] and, although the social behavior of *Eufriesea mexicana* is not known, the orchid bee tribe, Euglossini, to which both *Euglossa* and *Eufriesea* belong, is highly variable from solitary to communal to weakly social so we assign this taxon to the same category as *Euglossa dilemma. Lasioglossum albipes* is similar to *Euglossa dilemma* in its expression of a form of simple sociality that varies across the species [10] and *Ceratina calcarata*, though not representative of polymorphic social behavior, does exhibit the same type of weakly social behavior as the Euglossines and *L. albipes* [4], so is grouped in the same social category as these other taxa.

##### *1.3 Genome alignments*

Scaffolds shorter than 1,000 bases were first filtered from all genome assemblies. *De novo* repeat libraries were obtained for each genome using standard approaches as follows. We used RepeatModeler [11] to obtain initial repeat libraries and eliminate redundant repeat sequences of at least 80% similarity. We then filtered out any sequences that matched to known protein-coding sequences in UniProt and also to *Drosophila melanogaster* sequences using BLASTX. Sequences at least 50% similar across at least 50% of their length to any proteins were removed from the repeat libraries. We combined these *de novo* repeat libraries with the RepeatMasker libraries for Arthropoda downloaded on March 8, 2017 and used all of these repeat sequences together to mask genomes using RepeatMasker [12]. RepeatModeler repeatedly crashed on the genomes for both *A. mellifera* and *E. mexicana* so the repeat library from the closely related *Apis florea* was used to mask these sequences.

The resulting soft-masked genome sequences were aligned in all pairwise combinations using LAST v. 914 [13,14]. We used LAST-TRAIN with parameters "--revsym --matsym --gapsym -C 2 -E 0.01" to create alignment parameters for each pair of genomes. Alignments were performed using parameters "-m 100 -E 0.01 -C 2" and we used maf-swap to reduce the alignments to just the single best alignment for each region. We processed the resulting MAF files with code available here: https://github.com/berrubin/BeeGenomeAligner. First, we filtered out alignments that appeared to be spurious by examining synteny. We required that at least 5% of the bases aligned from a particular scaffold from species1 be aligned to a single scaffold in species2 for any of those alignments to be included. This substantially reduced the number of very short and likely spurious alignments. LAST produces a large number of alignments that are relatively close together, but due to a lack of intervening alignable sequence, splits these into multiple loci. We merged those loci within 500 bases of each other in the *Apis mellifera* genome, requiring that they be aligned on the same strand and scaffold. This again served to reduce the number of loci analyzed. Any alignments shorter than 1,000 bases at this stage were discarded. Coding sequences from the official gene sets were soft-masked from all genomes. MAF files from different species were then merged into multiple sequence alignments by identifying loci with overlapping coordinates in the *A. mellifera* genome. These sequences were combined from all species and then realigned to each other using FSA [15]. For the FSA alignment, we used the "--anchored" and "--exonerate" options to improve quality and speed of alignment of large regions. We also soft-masked non-coding sequence for these alignments and used the "--softmasked" option (coding sequence was unmasked at this stage).

We hard-masked coding sequence and then performed sliding window analysis across these alignments examining 500 base windows, iterating by 250 bases each step. Sequences were only included in windows when they were composed of at least 50% known nucleotides (excluding gaps and N's). Alignments were filtered with trimAl using the "automated1" option to remove poorly aligned regions [16]. Filtered alignments shorter than 250 bases were discarded.

##### *1.4 The distribution of non-coding evolutionary rates*

In addition to comparing NCARs with different rates of evolution in association with social evolution, we also identified those NCARs evolving exceptionally quickly or slowly overall without consideration for differences in rates in particular lineages. First, we reconstructed branch lengths for the phylogeny by concatenating all proteins with orthologs in all 12 species totaling 1,052,985 amino acids using RAxML [17] with the PROTGAMMAWAG model of substitution. For each NCAR, we standardized total branch length to the branch lengths obtained from this full protein dataset, first trimming those tips not present in the current region of interest. This standardization made NCARs comparable across loci, providing a metric of total evolutionary change for each individual locus. We then sorted this standardized total evolutionary change in each NCAR and examined the 100 fastest-evolving and slowest-evolving gene-associated NCARs corresponding roughly to the $3^{rd}$ and $97^{th}$ percentiles of the rate distribution.

##### *1.5 Relative rates tests*

We log-transformed branch lengths before calculating relative rates, used a weighted regression to calculate rates, and used a minimum branch length cutoff of 0.001, discarding branches shorter than this length. We only examined genes represented by at least nine different lineages and only calculated significance for those with at least three representatives of the focal lineages. RERconverge identifies both those genes evolving at a higher rate in the foreground branches as well as those evolving at a slower rate with Kendall correlation tests. We examined each of these groups separately, and included loci in downstream analyses with $p < 0.05$. Because the small sample sizes for tests of each individual gene mean that p-values have a relatively high minimum value, multiple-test correction was ineffective for this dataset and was not applied.

Using this methodology, we were able to test for an association between any degree of social behavior with 4,852 genes, and an association between evolutionary rates and complex sociality across 3,288 genes. GO terms were assigned to orthologous groups of genes using Trinotate [18] annotations of *Apis mellifera* representatives. We then identified enriched GO terms in sets of genes identified by RERconverge using GO-TermFinder [19].

Estimated branch lengths for NCARs were examined using RERconverge as was done for the coding regions.

##### *1.6 Gene tree discordance*

Gene tree discordance can cause erroneous identification of substitution rate increases [33]. We evaluated the impact of such specious characterizations on our results by identifying both coding sequences and NCARs with signatures of gene tree discordance. We used RAxML v7.3.0 [32] to reconstruct phylogenies for individual coding and non-coding loci. The model PROTGAMMAWAG was used for inferring phylogenies from coding sequences and GTRGAMMA was used for inferring phylogenies for non-coding loci. We then evaluated the fit of the sequence data to the species and gene trees using FastTree [34] and compared these inferred likelihoods using CONSEL [35].

#### *2. SUPPLEMENTARY RESULTS*

##### *2.1 Enrichment of sequence features in outlier NCARs*

Compared to the 3,233 gene-associated NCARs, the 100 fastest evolving gene-associated NCARs were significantly underenriched for upstream regions (hypergeometric test, p = 0.003, 0.26-fold underenrichment) and downstream regions (hypergeometric test, p = 0.03, 0.64-fold underenrichment) and over-enriched for 5'-UTRs (hypergeometric test, p = $3.1\times10^{-10}$, 4.5-fold enrichment). Promoters were significantly underenriched in the 100 slowest evolving NCARs (hypergeometric test, p = 0.022, 0.53-fold underenrichment) and downstream regions were overenriched (hypergeometric test, p = 0.032, 1.64-fold enrichment). Of the 255 miRNAs available from mirBase, nine overlapped with NCARs. None of these were in either the 100 fastest or slowest evolving NCARs.

##### *2.2 Leave-one-out validation of complex eusociality tests in RERconverge*

The RERconverge test for rate changes in taxa with complex eusociality (e.g. *Apis* + *Melipona*) yielded 240 NCARs with significantly faster rates of evolution and 237 NCARs with significantly slower rates. To assess how much of this signal is driven by *Apis* alone, we excluded *M. quadrifasciata* from these tests and find 242 NCARs have significantly faster rates and 223 have significantly slower rates. Of these *Apis*-specific NCARs, 154 faster and 131 slower regions overlap between the two tests. Nine GO terms are significantly enriched in those NCARs evolving significantly faster in complex eusocial taxa and only one of these is also enriched in those NCARs evolving more quickly in the three *Apis* lineages (GO:0000122; negative regulation of transcription from RNA polymerase II promoter; FDR-corrected p = 0.0098; Table S11). This is the only term enriched in the loci evolving faster in *Apis* lineages alone. However, eight of the nine GO terms showed significant signatures of enrichment before multiple test correction (Table S11), suggesting that a trend exists in the data even in the absence of *Melipona*. No GO terms were significantly enriched in slowly evolving regions, as is the case for the test of all complex eusocial lineages.

Similarly, we also excluded *A. mellifera* from these analyses and found 64 and 89 NCARs are evolving faster and slower in the remaining complex social species. Of these, only 43 and 45 overlap with the results from the test of all complex eusocial lineages. This suggests that *A. mellifera* may be driving a large part of the observed signal in the full dataset. However, three GO terms were significantly enriched in faster evolving NCARs in this test, one of which is also enriched in the full dataset test (GO:0009653; anatomical structure morphogenesis; FDR-corrected P = 0.018). The two other terms are GO:0010927 (cellular component assembly involved in morphogenesis; FDR-corrected p = 0.026) and GO:0007424 (open tracheal system development; FDR-corrected p = 0.050). Five of the nine GO terms identified as significant in the full test of complex eusocial taxa do show signatures of enrichment before multiple test correction (uncorrected $p < 0.05$; Table S11). Again, no GO terms were significantly enriched in slowly evolving regions.

Finally, excluding *A. florea* from the RERconverge analysis yields 79 genes evolving faster and 97 genes evolving slower in taxa with complex sociality with overlaps of 47 genes with the full dataset results in both categories. A single GO term was significantly enriched in the faster evolving NCARs (GO:0010927; cellular component assembly involved in morphogenesis; FDR-corrected p = 0.017; Table S11). This GO term is not present in the full dataset, though is related to several enriched terms. Again, five of the nine GO terms identified in the full test of complex eusocial taxa are significantly enriched before multiple test correction (uncorrected $p < 0.05$; Table S11) and no GO terms were significantly enriched in slowly evolving regions.

For each of these three tests, we focused in particular on the nine GO terms significantly enriched in the faster-evolving loci from the full dataset. The enrichment of these terms from the leave-one-out tests are given in Table S11. While few pass correction for multiple testing, the majority of GO terms identified as enriched in the full dataset do show some signature of enrichment. These results give us confidence that our results are not being driven by particular taxa.

Note that there are four lineages with complex sociality in the full dataset: *A. mellifera, A. florea, M. quadrifasciata*, and the lineage ancestral to the two *Apis* species. Thus, when *M. quadrifasciata* is excluded, that reduces the foreground lineages to three whereas when either of the *Apis* species is excluded, that reduces the number of foreground lineages to two. This more drastic reduction in number of lineages included in the test may explain the smaller number of regions identified. However, these results may indicate that the signal is driven largely by the *Apis* lineages and that the convergent behavioral evolution in *M. quadrifasciata* contributes little.

##### *2.3 Assessing the contribution of signal from* Melipona

We, therefore, also performed additional tests to assess the influence of *Melipona* on the results. Both *B. impatiens* and *B. terrestris* are equally related to *Apis* as *Melipona* but do not have complex social behavior so represent lineages for which we do not expect to see the same degree of convergent evolution. We performed two additional RERconverge runs which included the three *Apis* branches as well as the two *Bombus* species individually as the focal lineages. The degree of overlap between these tests and the test of all lineages with complex sociality demonstrates how much sequence-level convergence is present.

For the test including *Bombus impatiens* as a focal lineage with the three *Apis* lineages, 176 and 193 NCARs were found to be evolving significantly faster and slower, respectively. For *B. terrestris*, 188 and 172 NCARs were evolving faster and slower, respectively. Recall that 240 and 237 NCARs were found to be evolving significantly faster and slower in the test of all complex social lineages. While these numbers are not drastically different, they do suggest that there is more convergence in rates of evolution among all complex eusocial lineages than between the *Apis* lineages and either *Bombus* species. Only a single GO term is significantly enriched in the fast-evolving loci in *Apis* and *B. terrestris* (GO:0017124, SH3 domain binding; FDR-corrected $p = 0.042$). Similarly, only one GO term is significantly enriched in the fast-evolving loci in *Apis* and *B. impatiens* (GO:0003677, DNA binding; FDR-corrected $p = 0.011$). Neither of these GO terms were also found to be enriched among loci evolving faster in the test of all complex eusocial lineages. Thus, there appears to be greater convergence in molecular pathways among the complex eusocial *Apis* and *Melipona* than between *Apis* and *Bombus*.

##### *2.4 Null expectations for RERconverge and GO enrichment in NCARs*

Nine GO terms were enriched in the NCARs found to be evolving faster in the test of complex social taxa and these were of particular interest in our examination of the null expectation for GO enrichment. In the tests for GO enrichment across the 1,000 iterations of random foreground lineages, faster evolving NCARs were found to be enriched in GO terms enriched in complex eusocial lineages four times. Three of these were enrichments for GO:0030182 (neuron differentiation) and one was for GO:0000122 (negative regulation of transcription from RNA polymerase II promoter). In the slower-evolving NCARs, two iterations were again enriched for GO:0000122 (negative regulation of transcription from RNA polymerase II promoter) and two were enriched for GO:0045165 (cell fate commitment). Given this small degree of overlap in enriched GO terms between 1,000 tests of random foreground branches and complex eusocial lineages, it is clear that complex eusocial lineages do share signatures of rate changes and that the functional enrichment found is different than random expectations.

We also created 1,000 sets of 240 (the number of NCARs evolving significantly faster in complex eusocial taxa) random NCARs and examined the associated genes for GO enrichment. 219 of 1,314,513 (0.017%) GO terms tested were significantly enriched (FDR-corrected $p < 0.05$) including three GO terms that were also identified in tests of complex social lineages: GO:0001745 (compound eye morphogenesis), GO:0030182 (neuron differentiation), and

GO:0000122 (negative regulation of transcription from RNA polymerase II promoter). Thus, identifying each of the nine GO terms identified in the complex eusocial taxa at random is very unlikely (permutation test $p \leq 0.001$). Overall, the small number of overlapping GO terms found from random foreground lineages and the tests of complex social lineages suggests that these signals are not the result of random sampling.

##### *2.5 Motif turnover*

HOMER predicted the presence of the 147 sequence motifs 9,677 times in at least one species in those NCARs with significantly faster rates of evolution in complex species and 9,311 times in regions with significantly slower rates. We examined all motifs that occurred in any taxon within every NCAR for differences in frequency between socially complex and all other taxa in that NCAR. For example, an NCAR for which a particular motif occurred in the sequence of all three taxa with complex sociality and no other species would be identified as more abundant in taxa with complex sociality. Such significant differences ($\chi^2$ $p < 0.05$) in frequency between social types existed for 2,442 motif occurrences, 1,678 (69%) of which were more abundant in socially complex taxa and 764 (31%) of which were more abundant in all other taxa. Of these differentially present motif occurrences, 242 (10%) were in NCARs evolving significantly faster in socially complex taxa and 129 (5%) were in regions evolving significantly slower. However, of the 1,678 motif occurrences for which the motif was more frequent in taxa with complex sociality, 148 (9%) were in regions evolving significantly faster in this group, a significant underenrichment (hypergeometric test, $p = 0.005$) and 104 (6%) were in regions evolving significantly slower in these taxa, a significant overenrichment (hypergeometric test, $p = 0.001$). In motifs more abundant in all taxa that do not exhibit complex sociality, 94 (12%) were in regions with faster evolution in taxa with complex sociality, a significant overenrichment (hypergeometric test, $p = 0.005$) and 25 (3%) were in regions with slower evolution, a significant underenrichment (hypergeometric test, $p = 0.001$). Therefore, NCARs that evolve at faster rates in particular taxa appear to lose binding motifs in those taxa. This pattern suggests that faster evolution may be indicative of a loss of conserved binding motifs, rather than a gain of new binding motifs. However, newly emerged binding motifs may not be widespread enough to be identifiable by *de novo* motif discovery.

Many of the significant differences in frequency included the same motif sometimes present in greater abundance in complex eusocial taxa and sometimes present in greater abundance in solitary taxa, depending on the NCAR currently being examined. However, there were 20 unique motifs present at significantly higher frequencies only in regions evolving more rapidly in complex eusocial taxa, as well as 19 different motifs that occurred at higher frequencies only in regions evolving more slowly in complex eusocial taxa relative to all other species ($\chi^2$ test, $p < 0.05$; Table S14). Higher frequencies of these unique motifs suggest that there may be some convergence in the transcription factors co-opted as social behavior has become more elaborate in each lineage. Below, we also examined motif evolution in detail in those NCARs associated with genes involved in neuron fate commitment and neuron differentiation, finding several motifs involved in neural development associated with complex eusociality (SI 2.7, 2.8).

In general, there appeared to be rapid turnover of motif sequences with loss frequently followed by gain of a new motif. Among those NCARs where a motif was present in significantly fewer taxa with complex eusociality than in all other taxa, the numbers of motifs present at significantly greater frequency in these species was not significantly different than the number of motifs present at significantly greater frequency in other species (one sample Wilcoxon test $p = 0.09$; mean difference between number in complex eusocial lineages and in other lineages = -0.092). Therefore, it appeared that when ancestral motifs were lost, novel motifs were gained, perhaps to replace them.

***2.6 Motif abundance***

For each species, we counted the proportion of NCARs where each motif appeared at least once and tested for differences in these proportions between the species with complex sociality and all other taxa. Using this approach, we found 18 motifs that differ significantly in the number of NCARs in which they occur between taxa with complex sociality and all other species (Wilcoxon rank-sum test, $p < 0.05$; Table S13). Eight were significantly more abundant in complex social species and 10 were more abundant in other species. Four of these were also significantly different in abundance when correcting for phylogenetic history (PGLS test, $p < 0.05$; Table S13). Using the same approach, we also identified nine sequence motifs that differ significantly in the number of NCARs in which they occur between taxa with any degree of sociality and strictly solitary species (Wilcoxon rank-sum test, $p < 0.05$; Table S12). Seven of these were also significantly different using phylogenetically corrected tests (PGLS test, $p < 0.05$; Table S12).

***2.7 Neuron fate commitment motif turnover***

Several of the NCARs evolving significantly faster in taxa with complex sociality that were associated with genes involved in neuron fate commitment also showed patterns of motif gain and loss in these taxa (Table S10). There were two windows evolving at significantly different rates in lineages with complex sociality associated with *drk*, both of which were in the 3'-UTR. However, one of these windows was evolving significantly faster and one was evolving significantly slower. In the slower evolving NCAR, there were three sequence motifs that were present in all complex social lineages and only in one or none of the other lineages (i.e. all were present in significantly more lineages with complex sociality than other taxa ($P < 0.05$)). These motifs were VCTBAGGG, GYWVTCAY, and CCGTAAGCGCAT. The motif VCTBAGGG is particularly interesting as it was highly abundant across NCARs and was present in the sequence of species with complex sociality in significantly more NCARs (Table S13). This motif had a match score of 0.75 to a binding site for *AP-2α,* a gene involved in neurodevelopment. The motif GYWVTCAY was the $12^{th}$ most commonly encountered motif and had a 0.77 score to a binding motif for *Su(H)* which was associated with memory and the gene *Notch.* CCGTAAGCGCAT is the most abundant motif overall, was significantly more abundant in species lacking complex sociality, but had no clear known ortholog.

Four NCARs were evolving significantly faster in complex social lineages associated with the transcription factor *elB,* all of which were in the first intron of this gene. One NCAR had lost two sequence motifs in the complex lineages which were present in all other lineages (CAATCAGT and CTGACTAGTA). The first of these, CAATCAGT, had a similarity score of 0.85 to a *onecut* binding motif in *Drosophila*, another gene involved in nervous system function. The second, CTGACTAGTA, did not have a match with a score of at least 0.75 to any previously characterized motifs. A third motif in this NCAR, TAWNGTGCBG, was present in both *Apis* species and no other species. No similar motif in any other species is known. In the second NCAR, there were three motifs present in two species with complex sociality and absent otherwise (TAAGCACT, GGGCCCCG, and WTRGVGRG). The motif TAAGCACT has a 0.80 score to a binding site of *vvl* which is heavily involved in brain development, GGGCCCCG has a 0.78 score to a binding site of *Tcp* about which little is known, and WTRGVGRG has no match with a score of at least 0.75. No motifs occurred in significantly different numbers of taxa in the other two NCARs associated with *elB.*

A single region downstream of *babo* was evolving significantly faster in taxa with complex sociality. Two sequence motifs were present in all three of the complex eusocial species and absent in all others (GTWYHWWDTTTT and ATGTCACA). One, ATGTCACA, had a similarity score of 0.79 to an *hth* motif in Drosophila but the second, GTWYHWWDTTTT, did not have similarity score of at least 0.75 to any known motifs.

##### *2.8 Neuron differentiation motif evolution*

There were nine NCARs evolving significantly faster in taxa with complex eusociality that were associated with genes involved in neuron differentiation (and not neuron fate commitment) that showed patters of motif gain and loss across taxa (Table S10). For *tkv* (FBgn0003716), there was a single NCAR in the first intron evolving faster in taxa with complex eusociality. The motif TTATATAGTGAA was present in all three complex eusocial taxa in this NCAR and was not present in any other taxa. This motif did not have a match score of at least 0.75 to any known motifs.

Two NCARs upstream of *Arm* were both evolving significantly faster in taxa with complex eusociality. The first, 6,118 bases upstream, included three motifs present in a significantly greater fraction of taxa with complex eusociality than other taxa (GCCGGCYG, CCCTGCCT, and CTCCCTCC). GCCGGCYG had a 0.90 match score to the binding site of ethylene response factor 6 in *Arabidopsis* and was present in two taxa with complex eusociality and no others. CCCTGCCT had a 0.75 match score to a binding domain of the gene *HNRNPH2* in humans and was present in all three taxa with complex eusociality as well as *Lasioglossum albipes*. CTCCCTCC was present in all three complex eusocial taxa as well as *Eufriesea mexicana* and was a 0.83 match score to a binding site of the gene *B52* in *Drosophila,* a gene involved in gene expression regulation. Six motifs were more abundant in the second upstream NCAR, also including CTCCCTCC which was again present in all three complex eusocial taxa but no others. AGAGAGAGAGAG had a 0.88 match score to a binding site of *Trl*, a transcription factor involved in chromatin modification in *Drosophila,* and was present in all three complex taxa and no others. SDBCSYCTCT was present in *A. mellifera* and *M. quadrifasciata* but no others and had a 0.82

match score to a binding site of *SRSF10* in humans, a splicing factor. GCTCACATAG, MTCSCCCTCG, and KTATGGYMCW were each present in two complex eusocial taxa and no others and did not have match scores of at least 0.75 to any known motifs.

A single NCAR downstream of GB54569, which has a BLASTP hit to the nervous-system expressed gene *Gfrl* with an e-value of 0.7 but does not have a *Drosophila* ortholog in OrthoDB, had the motif ATAAATAG in the two *Apis* species and no other taxa. This motif had a 0.91 match score to a binding site of the *bin* transcription factor in *Drosophila.*

An NCAR in the last (third) intron of *dsx* was evolving significantly faster in taxa with complex eusociality and, in this case, three motifs had been lost in these taxa. These included GCCGGCYG, which was also gained in the a*rm* associated NCAR discussed above and was present in no taxa with complex eusociality and 7/8 other taxa. GCATAATGCC was present in *M. quadrifasciata* and all non-complex taxa and CCGGGCTA was absent in all complex eusocial taxa and present in 7/8 other taxa. Neither have a match score of at least 0.75 to any known motifs.

For *sub*, a single downstream region was evolving faster in species with complex eusociality and both *Apis* species have gained the motif WTRGVGRG which was not present in any other taxa. This motif did not have a match score of at least 0.75.

An NCAR in the first intron of the gene *sens* was evolving significantly faster in complex eusocial taxa. A single motif had been lost in these taxa and was present in 6/8 other taxa. This motif (GTGCGGCC) had a 0.75 match score of a binding site for *STP1* in yeast.

For *hh*, an NCAR in the first and only intron was evolving significantly faster in taxa with complex eusociality. Two motifs had been gained in these taxa, both of which were present in all three taxa with complex eusociality and only a single other taxon. These were WTRGVGRG which did not have a match score of at least 0.75 to any known motifs and TATTATCG which acts as a binding domain for the *Drosophila* gene *qkr58E-1,* of which little is known.

Two NCARs in the first intron of the gene *FoxP* were evolving faster in complex eusocial taxa, one of which had lost a motif in these taxa which is present in 6/8 other taxa. This motif (DGRCGSMYBN) did not have a match score of at least 0.75 to any known motifs.

For the gene *mbl*, a single NCAR in the first and only intron was evolving faster in complex eusocial taxa and a sequence motif (GTGCGGCC) which is present in 6/8 other taxa had been lost in taxa with complex eusociality. This is the same motif as was lost in the intron of the gene *sens* discussed above.

##### *2.9 Expression bias in foragers and nurses*

Of the 3,610 genes represented on the microarray used to compare queens and workers of *Apis mellifera* [20], 507 were proximal to one of the NCARs, 60 of which were significantly biased towards queens and 76 of which were biased towards workers. This represented a significant 1.52-fold under representation of the queen-biased genes (hypergeometric test, $p = 0.00003$). Worker-biased genes were represented at the expected frequency (hypergeometric test, $p = 0.22$, 1.08-fold under enrichment). Queen- and worker- biased genes were not over or under enriched in the 100 fastest or slowest evolving regions.

Worker-biased genes were significantly under enriched 2.17-fold (hypergeometric test, $p = 0.043$) in regions evolving significantly slower in species with complex sociality. No other tests of enrichment between fast and slow-evolving regions in association with social evolution were significant. There was some non-significant overlap between genes that were biased in expression and NCARs evolving at significantly different rates in RER tests and those detected as outliers either on the fast or slow end (Table S15).

Comparisons of genes differentially expressed in foragers and nurses also yielded equivocal results. There were 1,040 unique genes represented on the microarrays used to identify genes differentially expressed in foragers and nurses of *Apis mellifera* [21] proximal to an NCAR. Of these, 113 and 48 were evolving significantly slower and faster in taxa with complex sociality. Of the 80 that were biased in expression in at least one behavior, 11 were evolving significantly more slowly in taxa with complex sociality and 5 were evolving significantly faster. We, therefore, did not find a signature of over or under enrichment in genes with differential expression associated with NCARs evolving at different rates in taxa with complex sociality (hypergeometric test, $p > 0.05$).

There were 39 NCAR-associated genes biased in expression towards nurses and 59 were biased in expression towards foragers. These represented a significant 1.26 -fold enrichment of forager-biased genes (hypergeometric test, $p = 0.035$) and a 1.28-fold under enrichment in nurse-biased genes (hypergeometric test, $p = 0.049$). Nurse-biased genes were over enriched 2.28-fold (hypergeometric test, $p = 0.028$) in the 100 fastest evolving NCARs. We found no significant under or over enrichment in the 100 slowest evolving NCARs.

Of the 15,314 genes in the *A. mellifera* OGS v3.2, 209 were found to be queen-biased and 276 were found to be worker-biased in four day old larvae [22]. Only 16 and 31 NCARs were proximal to queen- and worker-biased genes, respectively. The NCAR proximal to one of these genes was detected as evolving significantly faster in complex eusocial taxa (GB53274). There was no other overlap between genes with caste-biased expression in larvae and significant differences in evolutionary rates indicating no significant overlap between these datasets.

***2.10 Overlap of genes and GO terms with previous studies***

Coding sequence evolution was previously correlated with social evolution in 10 of the 12 taxa examined here [3]. This previous study identified significant correlations between dN/dS ratios and level of social complexity, a more sophisticated approach than what we used in this study, meaning that we do not necessarily expect the same genes to be implicated.

There were 2,389 genes that were tested both in the previous study using dN/dS ratios and in the present study using the relative rates test to identify signatures of selection in taxa with complex sociality. In the previous study, 31 of these were identified as experiencing positive selection, 20 showed signatures of relaxation, and 41 showed signatures of purifying selection. Using the relative rates test, we find 169 genes with faster rates of change in taxa with complex eusociality (indicative of either positive or relaxed selection) and 159 with slower rates of change in these taxa. 13 of the genes identified using dN/dS as subject to positive selection and two of the genes identified as subject to relaxed selection were also identified as evolving faster in taxa with complex sociality using the relative rates test. Of the genes with dN/dS ratios indicative of purifying selection, 10 were also found to be evolving significantly more slowly using the relative rates test. While these numbers show significant overlap between the two types of analyses and datasets, there are clearly differences. Most noticeably, the relative rates test yields a much larger gene set implicated in selective processes associated with social evolution.

Only a single GO term was enriched in both our analyses of coding sequences and those presented previously [3]. This previous work showed enrichment of 14, 22, and 21 GO terms in the gene sets associated with positive selection, relaxed selection, and negative selection correlated with social evolution in these taxa. Using the relative rates tests here, we find three GO terms associated with faster evolving genes associated with complex sociality and 11 GO terms associated with slower evolving genes. A single GO term overlaps in the set of slower evolving genes and those previously associated with negative selection: GO:0008565, protein transporter activity. When examining the genes associated with all obligately social taxa, 25 and four GO terms are enriched in genes evolving slower and faster, respectively. Again, the same GO term is the only overlapping term with previous studies. Only two GO terms are enriched in the set of genes evolving faster in taxa with any degree of eusociality and in the set of genes evolving slower using the relative rates test. These two GO terms do not overlap with the results from the previous study.

No GO terms overlap between the non-coding analyses in this study and those previously found to be enriched in coding sequences.

##### *2.11 Little effect of gene tree discordance on detected changes in substitution rate*

Among coding sequences, 4,495 of 4,946 (91%) examined genes did not show significant support for an alternative phylogeny. Among those 3,288 genes included in the relative rates test, 318 (9.7%) were discordant. 257 genes were identified as evolving significantly faster in taxa with complex sociality, 27 (10.5%) of which showed significant evidence of gene tree discordance. 207 were identified as evolving significantly slower, 22 of which showed signatures of discordance (10.6%). We don't see any evidence for enrichment of loci with discordant gene trees being over represented among loci identified as evolving at significantly different rates (hypergeometric test, $p > 0.05$).

We checked 4,611 NCARs for signatures of alternative phylogenies and 335 (7.3%) of these showed significance. Of the 4,287 NCARs included in the RER tests, 302 (7.1%) were identified as having likely discordant evolutionary histories. Of the 237 NCARs identified as evolving significantly more slowly in taxa with complex sociality, six had discordant evolutionary histories and of the 240 NCARs evolving significantly faster, 21 (8.8%) had discordant phylogenies. This indicates a significant 2.78-fold under enrichment of slowly-evolving loci with discordant evolutionary histories (hypergeometric test, $p = 0.0016$). The faster-evolving loci did not show signatures of over or under enrichment (hypergeometric test, $p > 0.05$).

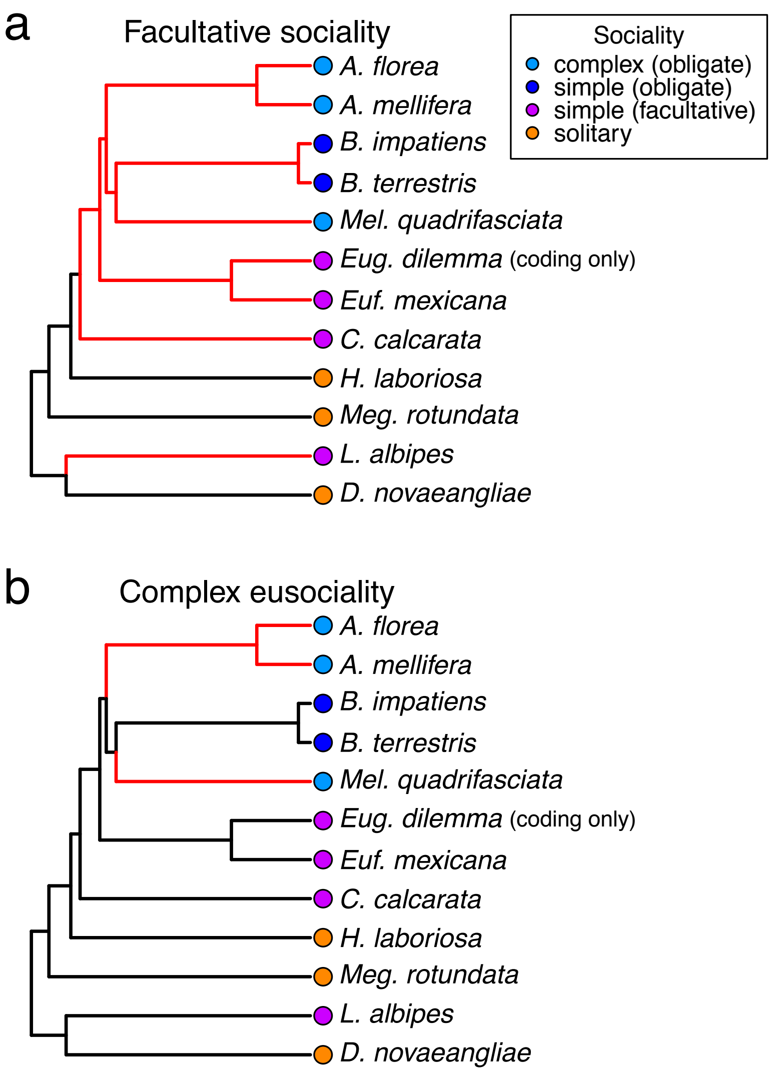


**Figure S1.** Phylogenies used to conduct relative rates tests with focal lineages colored in red.

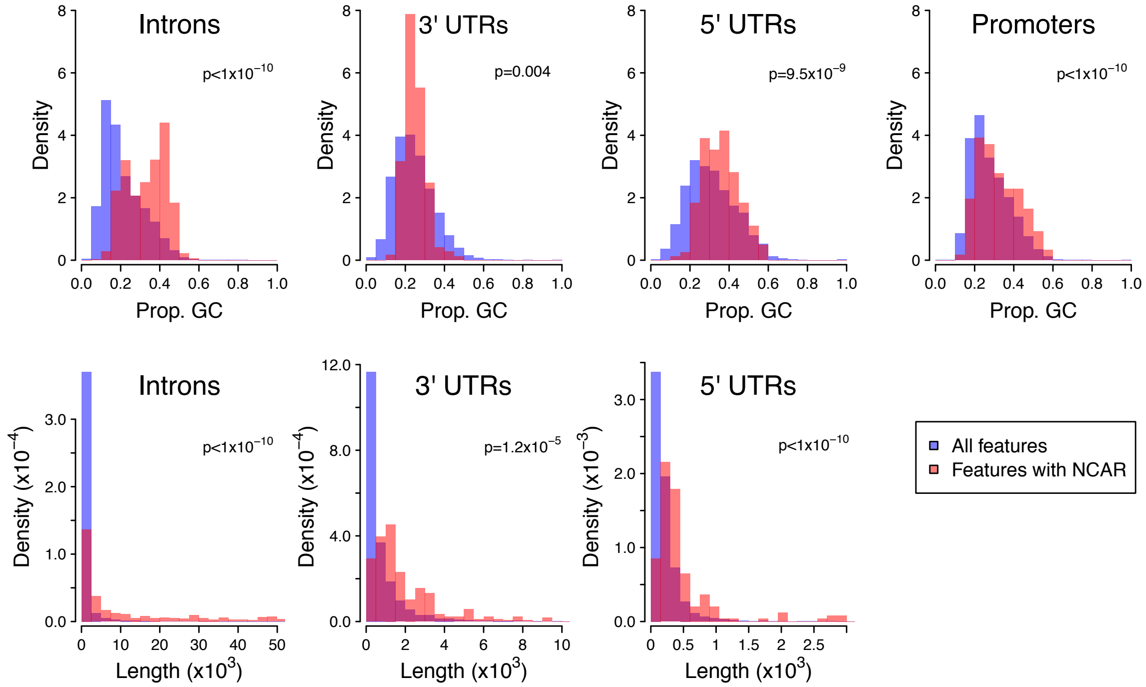


**Figure S2.** Distributions of GC-content and lengths of sequence features in which NCARs were identified (red) and all sequence features (blue) in the *A. mellifera* genome. P-values are the result of Wilcoxon rank-sum tests comparing these distributions. The length distribution of promoters is not shown because promoter length was fixed at 1.5kb.

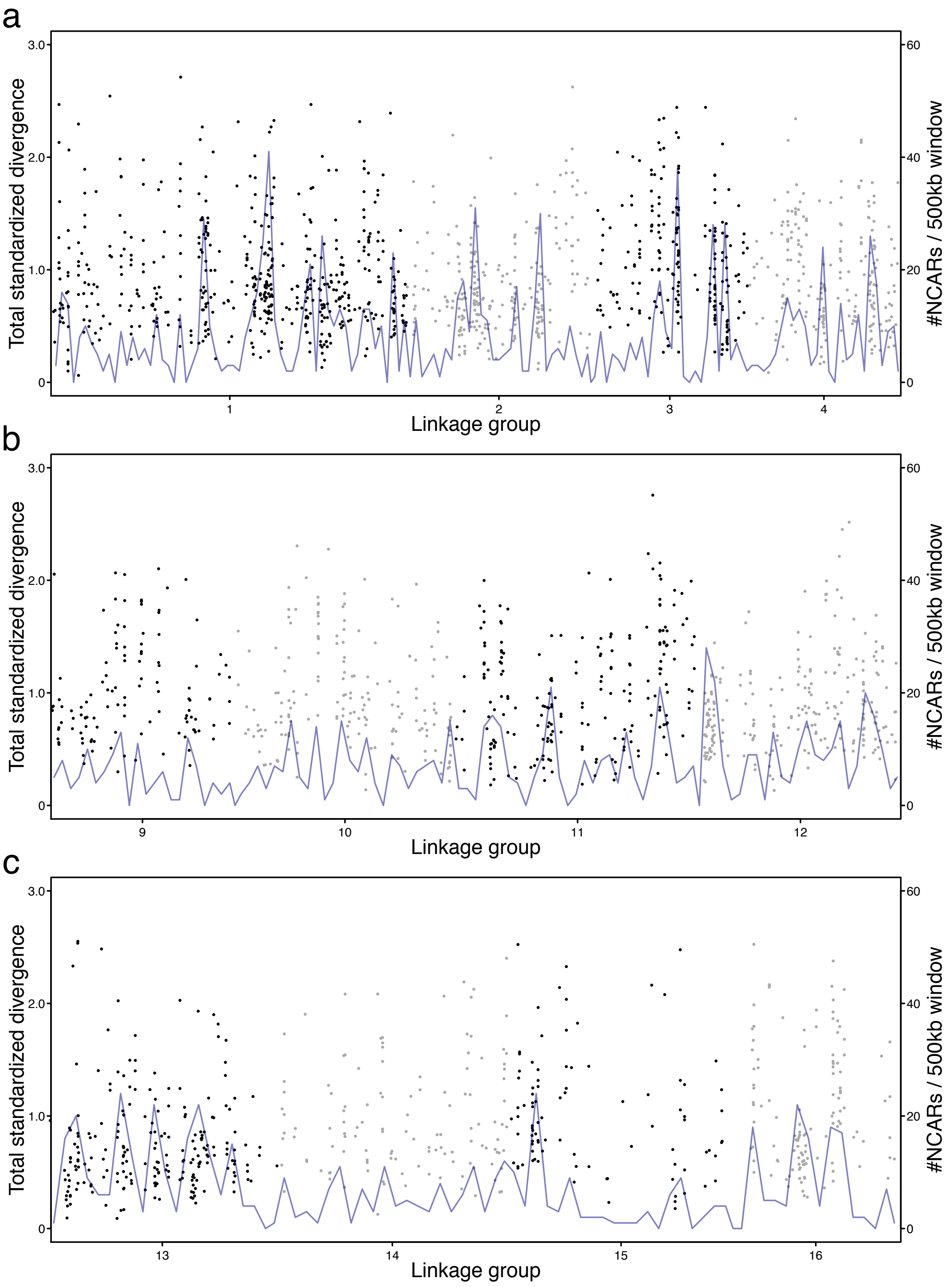


**Figure S3.** NCAR distribution across *A. mellifera* linkage groups 1-4 and 9-16 are represented as in Fig. 2b. Dots show the locations of NCARs. Black and gray colors are used to denote the linkage groups and the y-axis signifies a standardized measure of divergence for each region (detailed in methods). The blue line denotes the # NCARs present in each 500kb window.

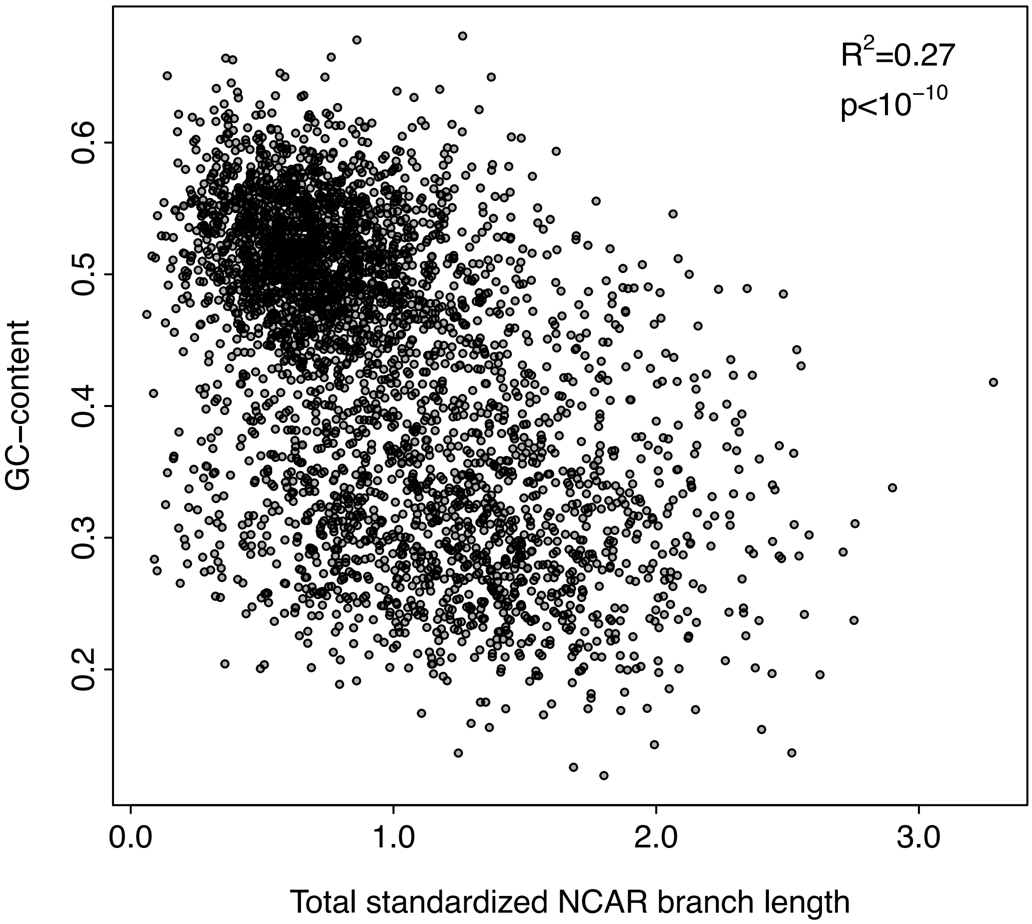


**Figure S4.** GC-content of *A. mellifera* sequence in each NCAR as a function of standardized total branch length of all taxa present in the NCAR.

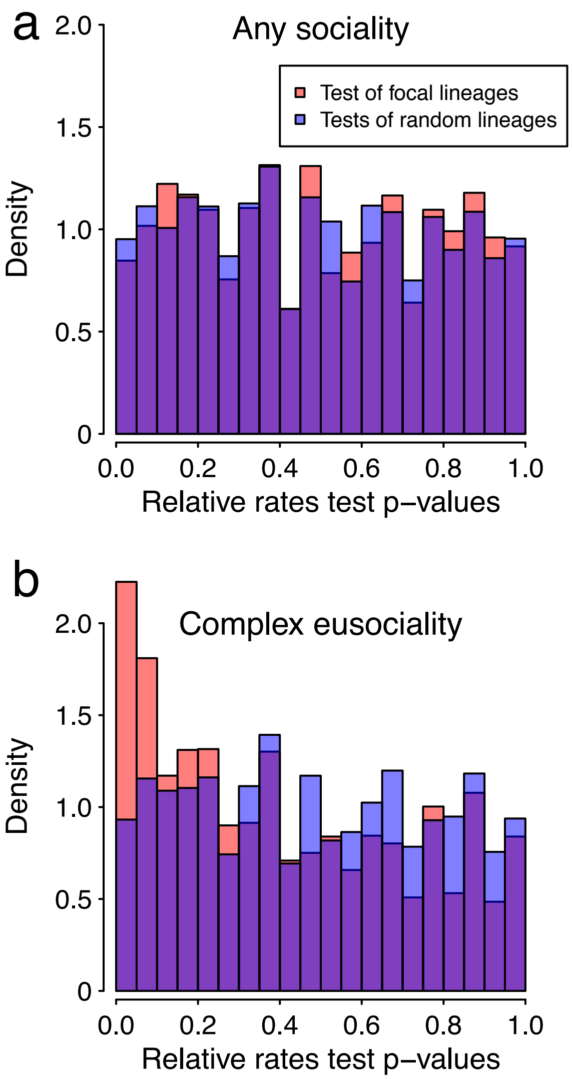


**Figure S5.** Distribution of p-values obtained from relative rates test including all lineages with any degree of sociality as focal taxa (a) and from relative rates test focused on only those lineages with complex eusocial behavior (b). Red bars show the results from the test of the indicated focal lineages and blue bars show the p-values obtained from 1,000 iterations of relative rates tests on randomly chosen focal lineages.

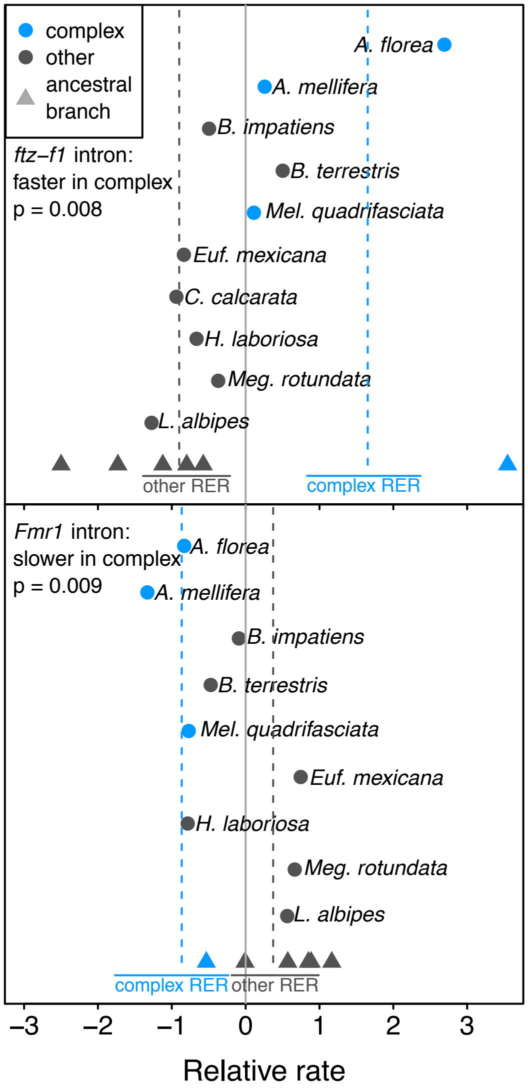


**Figure S6.** Two intronic NCARs associated with complex social behavior are key regulators of mushroom body neuronal remodeling (*ftz-f1*; [23]) and development (*Fmr1*; [24]). *ftz-f1* shows accelerated rates of change on complex social branches relative to the remaining branches in the tree (relative rates test, p=0.008). *Fmr1* shows significantly slower evolution on complex social branches (relative rates test, p=0.009).

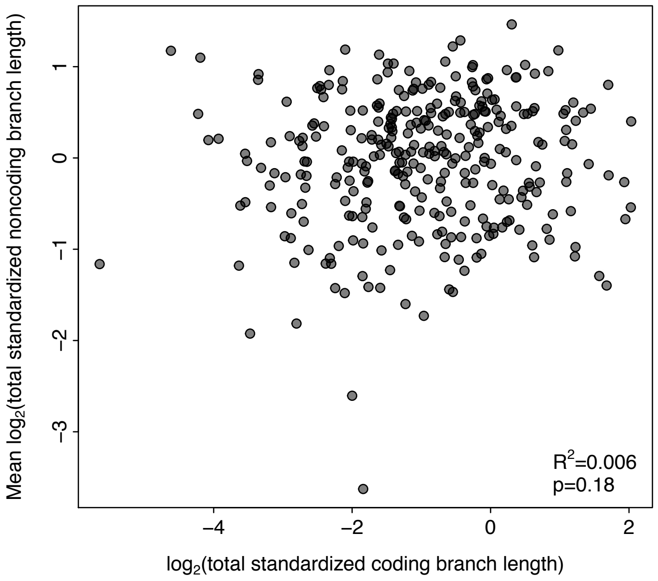


**Figure S7.** Log-transformed total branch length of coding sequences and proximal NCARs standardized to the branch lengths inferred from all a concatenation of all protein sequences. When multiple NCARs were associated with individual genes, mean standardized branch lengths were used.

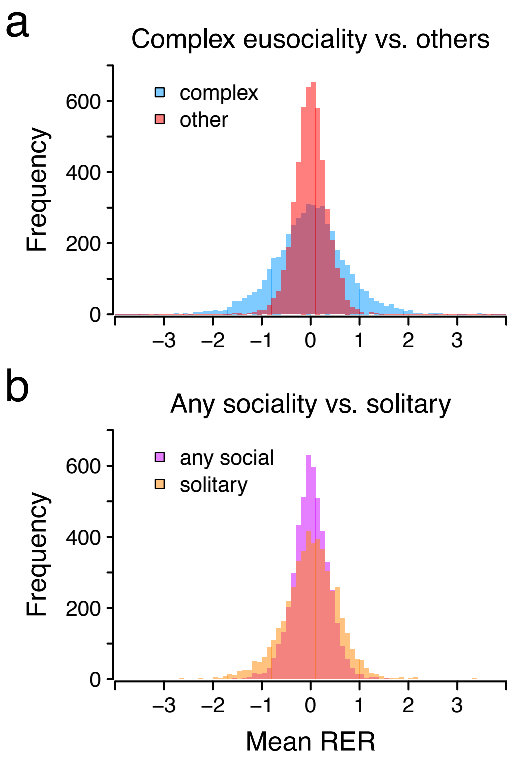


**Figure S8.** (a) Distribution of mean relative rates among taxa with complex sociality vs. others in all coding sequences. (b) Distribution of mean relative rates among taxa with any degree of sociality vs. strictly solitary taxa in all coding sequences.

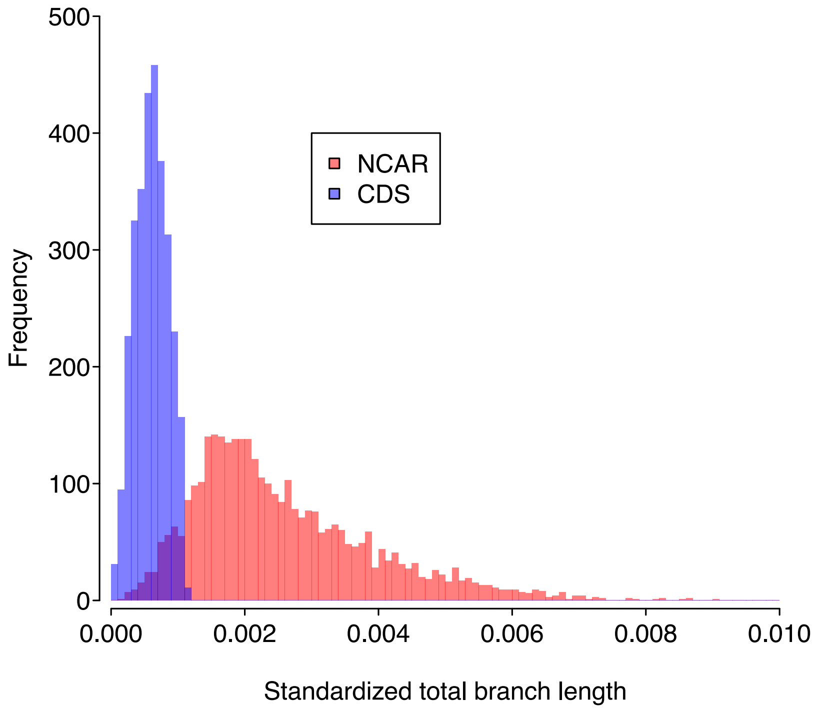


**Figure S9.** Distribution of total evolutionary change in all CDS's and NCARs analyzed. To make these measures comparable across loci and sequence classes, the standardized total evolutionary change was additionally divided by the number of bases in each locus.
